# Evaluation of Bacterial Microcompartment Cofactor Recycling and Permeability with a Model Guided *In Vitro* Assay

**DOI:** 10.1101/2025.08.20.671389

**Authors:** Brett Palmero, Christina Catlett, Charlotte Abrahamson, Sasha Shirman, Carolyn Mills, Michael C. Jewett, Niall M. Mangan, Danielle Tullman-Ercek

**Author notes:** Denotes equal contribution (Brett Palmero, Christina Catlett).

## Abstract

Biomanufacturing is a promising strategy for sustainable chemical production. However, challenges such as cofactor competition and low pathway flux prevent competitive titers. Some bacteria address these challenges by encapsulating metabolic pathways in bacterial microcompartments (MCPs), many of which contain dedicated cofactor recycling enzymes. We sought to determine how pathway cofactor recycling and intermediate sequestration in MCPs benefit pathway performance using an *in vitro* assay and kinetic model of the 1,2-propanediol utilization (Pdu) system. Guided by model simulations, we performed experimental design to characterize permeability, a key and difficult-to-measure property of MCPs. Using our model and measurements of metabolite concentrations over time, we estimate MCP permeability values in the range of 10^−5^ cm/s. We also demonstrated that NAD+/NADH recycling in the Pdu MCP benefits increased pathway flux. This study integrates experiments and systems modeling to advance our understanding of why pathways are encapsulated and to inform bioengineering applications.

## Introduction

Microbes are capable of producing a large variety of products from cheap and renewable sources, and substantial effort has been devoted to their use in sustainable commodity and specialty chemical production^1–7^. However, there are many roadblocks limiting pathway yield, including cofactor imbalances, buildup of toxic intermediates, and poor pathway kinetics^8–10^. Native metabolic pathways face similar roadblocks and have evolved multiple strategies to address them. One such strategy is to utilize spatial organization by sequestering pathways in proteinaceous polyhedral organelles called bacterial microcompartments (MCPs)^11^. Metabolosomes are a category of MCP that encapsulates the enzymes responsible for metabolizing niche carbon sources such as choline, ethanolamine, and 1,2-propanediol (1,2-PD) ^12–14^. MCPs are hypothesized to help pathway performance by increasing the local concentration of enzymes and substrates, creating private cofactor pools, diverting flux to the encapsulated pathway, and sequestering toxic intermediates^15–22^. However, it is unclear to what extent each of these benefits impacts metabolosome pathway behavior, or how the permeability of the shell influences pathway performance. Such an understanding would inform the selection of metabolic pathways of interest that may benefit from encapsulation and potentially increase biomanufacturing titers.

The *Salmonella enterica* subsp. Typhimurium 1,2-PD utilization (Pdu) MCP is the most well studied metabolosome, with tools to manipulate and investigate it, making it an excellent system for exploring the benefits of encapsulating pathways^21,23–32^. The Pdu MCP contains the enzymes required for the degradation of 1,2-PD into propionate for use as a carbon source (Figure 1A). In the Pdu pathway, the PduCDE catalyzes the conversion of 1,2-propanediol to propionaldehyde, a toxic intermediate. Several *in vivo* studies have demonstrated how sequestering propionaldehyde in the Pdu MCP helps protect the cell from harmful effects ^19,30^. Propionaldehyde is converted into propionyl-CoA and 1-propanol by the enzymes PduP and PduQ, respectively, which are coupled by NAD+/NADH recycling^16^. Propionyl-CoA is further converted into propionyl-PO_4_^2−^ by PduL. Propionyl-PO_4_^2−^ is converted by PduW into propionate. The initial step catalyzed by PduCDE proceeds faster than the steps catalyzed by PduP and PduQ, resulting in a kinetic imbalance. Encapsulating the Pdu pathway in an MCP is hypothesized to both sequester propionaldehyde and enhance pathway kinetics.

**Figure 1:**
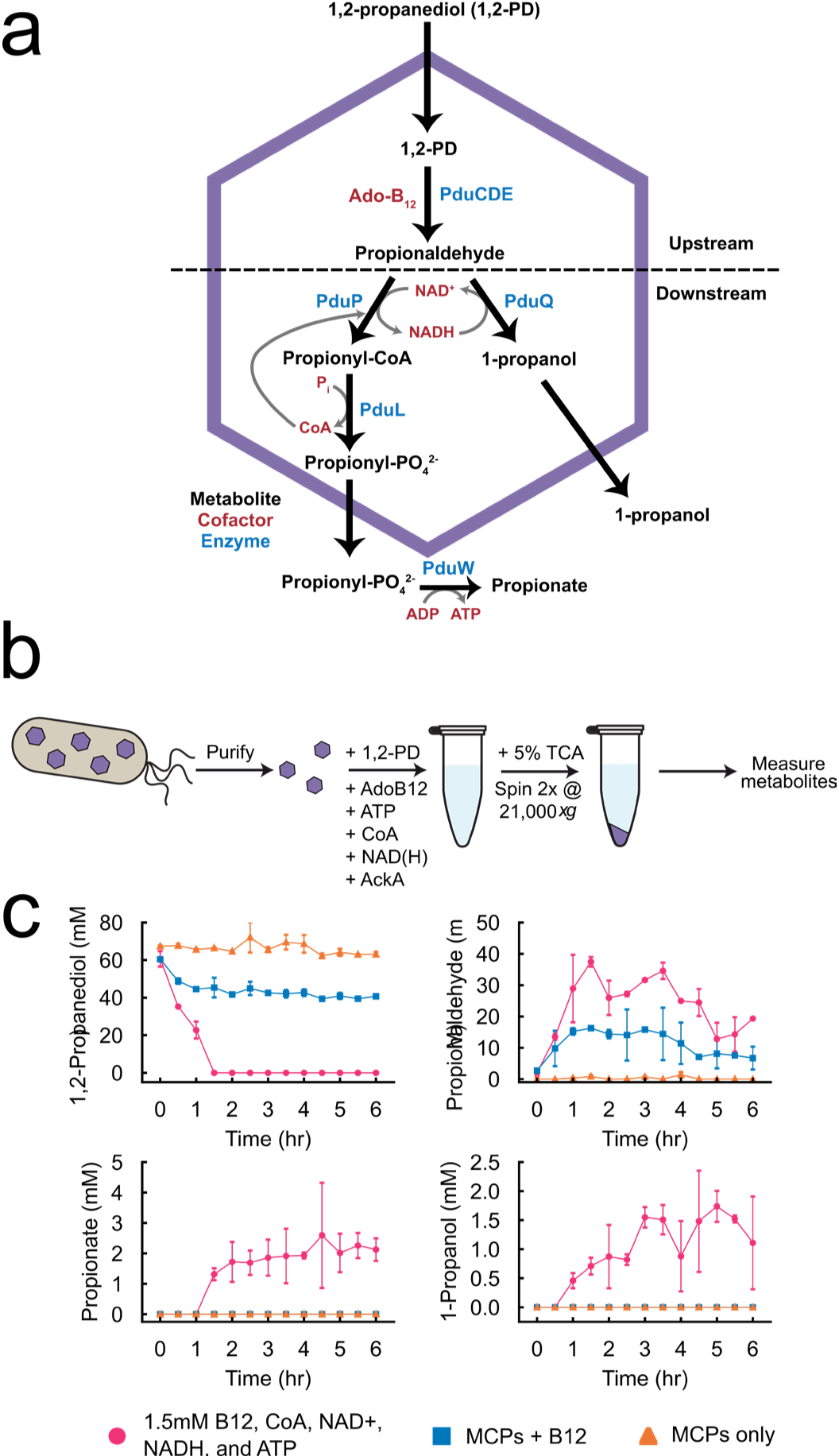
Pdu MCP function can be evaluated in vitro. **(A)** Diagram of 1,2-PD degradation pathway. **(B)** Schematic of the in vitro assay using purified MCPs. **(C)** Evaluation of wild type Pdu MCPs in vitro. Concentrations of Pdu metabolites were measured by HPLC. Error bars indicate standard deviation across three technical replicates.

To test hypotheses about how encapsulation in an MCP impacts pathway kinetics, assays that decouple flux from fitness must be developed. Altered pathway kinetics *in vivo* can result in growth defects, which can confound observed changes to pathway performance. Decoupling fitness and flux can be done in *in vitro* assays using purified Pdu MCPs in more controlled environments. For example, Cheng *et al.* assayed purified Pdu MCPs *in vitro* to determine if NADH could be recycled into NAD+^16^. The purified MCPs produced 62.3 μM propionyl-CoA, the product of PduP, when supplied with only 40 µM of NAD+, indicating recycling of NADH to NAD+. *In vitro* assays have also been used to verify the encapsulation efficiency of enzymes in the Pdu MCP by correlating decreases in observed activity with lower encapsulation efficiency^33^. However, each of these studies involving Pdu MCP *in vitro* assays has primarily focused on assessing one metabolite at a time. Concurrent quantification of multiple metabolites would yield a more complete picture of pathway performance *in vitro*.

Mathematical modeling can be used to infer biophysical properties that are difficult to measure directly. For instance, Young *et al.* used modeling to estimate the permeability of a programmable MCP shell to ATP^34^. Molecular dynamics simulations have also been used to estimate the permeability of synthetic MCPs^35^. By separating the internal and external dynamics of MCPs, modeling provides granularity not available in experiments. Given key parameters governing pathway kinetics and permeability, models can predict spatial and temporal dynamics of metabolite concentrations. However, these parameters are often unknown or poorly constrained experimentally. Prior studies have employed simulations spanning multiple orders of magnitude in parameter space to characterize the operating regimes of MCP systems, including the Pdu MCP and the carboxysome ^20,21,36^. These analyses demonstrate that shell permeability to different metabolites strongly influences the efficacy of encapsulation, affecting the degree of biochemical segregation of the MCP lumen from the cytosol and enhancement of pathway flux. Such work suggests an optimal range for permeability relative to the encapsulated pathway kinetics: if too high, metabolite sequestration is compromised; if too low, entry of substrate becomes overly rate-limiting.

Current attempts to quantify MCP shell permeability and metabolite sequestration remain primarily theoretical. In the absence of direct measurement, an assay pairing *in vitro* time-series data of metabolite concentrations with parameter estimation in a kinetic model could improve our certainty of permeability estimates. However, any such method must first overcome the expectation that parameter estimation for large systems with partially observed states and sparse data does not reliably converge to a unique, well-defined optimum^37^. This challenge motivates the need for experimental designs that constrain estimates of designated parameters-of-interest as much as possible. As new compartmentalized systems are characterized, the development of reliable assays to estimate key properties such as shell permeability can inform engineering design to increase pathway flux. In this context, model-guided experimental design offers a powerful approach for collecting data that constrains model parameters.

In this study, we developed a combined *in vitro* experimental and systems modeling approach to study the function of Pdu MCPs. We used this method to evaluate the hypothesis that one of the major benefits of encapsulating metabolic pathways inside MCPs is the creation of internal cofactor pools. The systems model was used to perform experimental design and determine the dynamic features and time points that will best inform parameter estimates of MCP shell permeability. A number of MCP enzyme and reaction environment configurations were investigated to isolate the effect of NAD+/NADH recycling on Pdu MCP function. Surprisingly, we found that cofactor recycling is beneficial but not necessary for Pdu pathway flux *in vitro*, implying the Pdu MCP can utilize both internal and external cofactor pools. We also found that cofactor recycling is vital to maintaining propionate production in cofactor-imbalanced environments. In addition, using the model and time-series data, we find a well-constrained estimate of MCP shell permeability consistent with previous predictions. Together, these results demonstrate the ability of *in vitro* whole-MCP assays to complement existing methods and show that internal cofactor recycling can improve MCP function *in vitro*.

## Results

### Development of a fully purified in vitro MCP activity assay

We sought to create an *in vitro* Pdu MCP system that allows for the evaluation of MCP activity *in vitro* by measuring the major metabolites of the Pdu pathway. Such a system also allows control over the reaction environment, for example pH, salt, or cofactor concentrations, which is not possible *in vivo,* and decouples pathway activity from cell fitness. To create our *in vitro* system, we combined purified Pdu MCPs with a reaction buffer containing the Pdu substrate 1,2-PD and the respective cofactors of the Pdu pathway (Figure 1B). Purification of intact Pdu MCPs was verified using transmission electron microscopy (Figure SI1). Since PduW is not encapsulated in the MCPs, we also added purified AckA, an acetate kinase from *Escherichia coli*, to convert propionyl-PO_4_ to propionate. We then sampled over time for metabolite analysis via high-performance liquid chromatography (HPLC). When we added all the required cofactors and AckA, we observed the production of propionaldehyde, propionate, and 1-propanol, indicating each Pdu pathway enzyme is active in this *in vitro* context (Figure SI2).

These initial *in vitro* experiments did not fully resolve the dynamics of the active Pdu pathway, because with these assays, we could not distinguish the metabolite concentrations interior and exterior to the Pdu MCP lumen. To overcome this limitation and access system dynamics relevant for estimating shell permeability, we employed kinetic-model-guided experimental design. We developed and applied a novel method to identify timepoints that would maximize information gain about pathway activity, given a kinetic model and known duration of experiment to reach equilibrium. This analysis revealed the peak in propionaldehyde concentration within the first five hours as the most informative dynamical feature of the data, prompting us to sample at 30-minute intervals for the first six hours of these experiments (Figure 1C).

Having observed activation of the Pdu pathway enzymes with added external cofactors, we next sought to determine if cofactor addition to this assay was necessary for activity. Currently, cofactors are hypothesized to enter the Pdu MCP upon assembly. Since only small amounts of cofactors were likely copurified with MCPs, we hypothesized that the addition of external cofactors might increase pathway flux. However, if ample amounts of cofactors were copurified, additional cofactor supplementation may not be necessary. To test this hypothesis, we used a simplified assay where we added purified Pdu MCPs and only 1,2-PD. In this assay, there was no observed consumption of 1,2-PD nor production of propionaldehyde, 1-propanol, or propionate (Figure 1C). We hypothesized that activity could be restored by the addition of coenzyme-B_12_ (B_12_) to catalyze the first step in the pathway. The B_12_ cofactor is not present when we purify Pdu MCPs because B_12_ synthesis is not active in the aerobic conditions in which we grow *S. enterica* for MCP purification. We repeated the experiment, this time with the addition of B_12_ to our base condition. This change conferred propionaldehyde production, confirming the activity of the first step of the reaction. This implies that B_12_ can diffuse and be utilized by the compartments when it is added exterior to the compartments in the reaction environment. However, we did not observe the products propionate or 1-propanol without the downstream cofactors. Once all of the cofactors were added, full pathway activation was achieved again. The necessity for cofactors in this process suggests that added cofactors are able to enter the lumen of the Pdu MCP shell and be utilized by the encapsulated Pdu enzymes. In addition, it indicates that if there is cofactor purified with the Pdu MCP, it is not sufficient for pathway activity.

### Experimental design guided by a systems model of the in vitro MCP system

We combined this *in vitro* Pdu MCP system assay with a kinetic model to guide sampling times and to provide the best estimate of the MCP shell permeability from indirect measurements. Briefly, our system-level model extends previous work describing MCP dynamics using a reaction-diffusion framework^21^. We assume a static population of identical MCPs and well-mixed conditions within both the MCP lumen and surrounding media, leading to a two-compartment model. The time evolution of metabolite concentrations (1,2-propanediol, propionaldehyde, propionyl-CoA, propionyl-PO_4_^2−^, 1-propanol, and propionate) as well as NAD+/NADH cofactor concentrations, is described by a system of ordinary differential equations. Importantly, unlike in our prior work, we explicitly model the PduP, PduQ, PduL and PduW/AckA mediated reactions and the NAD+/NADH cofactors. Other co-factors including ATP, B_12_, and CoA are assumed to be in excess and, thus, are not modeled explicitly.

Enzyme-catalyzed reactions follow reversible Michaelis–Menten kinetics, with reversibility constrained by free energy values obtained from literature, an update from previous models which assumed irreversible reactions^38^. Reasonable parameter ranges for kinetic rates and Pdu MCP geometry are based on imaging or drawn from existing literature. Transport between the two model compartments (MCP and media) is modeled by Fickian diffusion, with two distinct permeabilities, one for cofactors NAD+/NADH and one for the modeled metabolites. This distinction accounts for observed differences in small and large molecule diffusion through the pores in the MCP shell. Simulations are initialized with nonzero concentrations of 1,2-propanediol, propionaldehyde, NADH, and NAD+ inside and outside of each MCP, with the proportion inside to outside as an unknown parameter. Unavoidable variation in pipetting time will lead to variation in the internal proportion. For samples with a longer time between MCP addition and the initial sampling, a larger portion of the metabolites will have diffused into the MCP. Allowing initial nonzero concentrations of cofactors and upstream metabolites to exist in the MCP interior allows us to capture the uncertainty in timing.

To design experimental sampling, we simulated the system across a wide range of unknown parameters, including MCP permeability and enzyme kinetic rates. Rather than evaluating dynamics strictly as a function of time, we considered the measured metabolite concentrations as a function of the percentage of reaction progress toward steady state. We then calculated the sensitivity of metabolite concentrations to the varied parameters at each percentage of reaction progress using Principal Component Analysis (PCA)^39^. Evaluating the sensitivity to parameters in reaction progress space rather than time decouples the influence of dynamical trajectory shape from system time scale, allowing identification of informative features such as peaks, inflection points, or plateaus. Our analysis revealed that variation in MCP permeability most strongly affects the amplitude of the propionaldehyde peak, indicating that resolving this peak would best constrain estimates of permeability.

As the reaction progresses, the metabolite concentrations are initially relatively insensitive to variation in the parameters, peak in sensitivity between 60-80% progress, and then decrease in sensitivity; this sensitivity is captured by the magnitude of the first principal component of the PCA (Figure 2). Inflection points and peaks provide the most discriminative information between possible parameter sets. The contribution of different metabolites to the first principal component also changes over reaction progress. The magnitude of the first principle component indicates the information that can be gained by measuring the contributing metabolites at each point, as a measurement at those points will most drastically reduce the uncertainty in the parameters. Early in the reaction, the concentration of the upstream metabolites contribute most of the variance, with 1,2-PD dominating initially despite low absolute variation in comparison to the peak in propionaldehyde concentration. As the reaction proceeds, propionaldehyde dominates during its peak, and downstream products only become relevant in the final stage of the reaction, though overall variation is small in comparison to the variation at the propionaldehyde peak. As mentioned above, we used this information to inform our choice to take measurements at 30-minute intervals for 6 hours to resolve the peak in propionaldehyde concentration. While the decomposition of variation by metabolite in Figure 2 suggests that each metabolite need not be measured at each sample time, we simplified the optimal design as the HPLC detects all metabolites simultaneously and measured all metabolites at all recommended times. Finally, in order to map time to reaction progress, it is necessary to obtain an estimate or measurement of the final steady-state of the system. To compute progress toward equilibrium from our experimental data, we therefore collected a sample at t=24 hours where previous experiments found the system to have reached an approximate steady state (Figure SI3).

**Figure 2:**
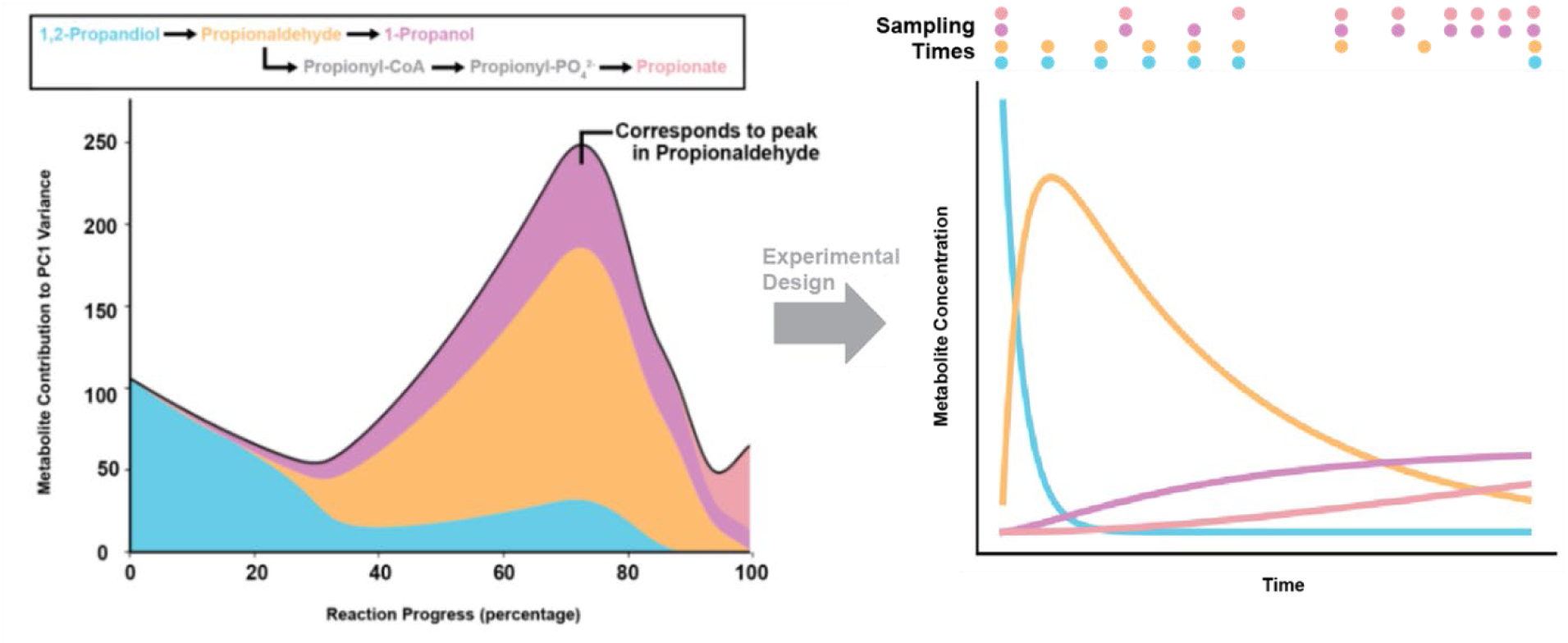
Mapping from time to progress coordinates allows for comparison of reactions on differing timescales. Decomposition of variation performed via Principal Component Analysis (PCA) determines measured state sensitivities to parameters, resulting in recommendations for sampling times. This early-reaction sensitivity is dominated by 1,2 Propanediol and Propionaldehyde, while the later reaction sensitivity was dominated by Propionyl-CoA and Proprionate. To capture the propionaldehyde peak, the first 6 hours were sampled at 30 minute intervals, and a final 24 hour time point.

### NADH recycling is beneficial but not required for Pdu pathway function

One of the major hypothesized benefits of MCPs is the presence of private cofactor pools that are internally recycled within the MCP. In Pdu MCPs, PduP and PduQ are hypothesized to be coupled by NAD+ and NADH recycling, as these cofactors are required for 1,2-PD degradation pathway^16^. While PduP is directly necessary to produce propionate for ATP generation and carbon utilization, PduQ is hypothesized to help maintain the NAD+ pool for PduP but is not directly responsible for bacterial growth on 1,2-PD. In fact, the deletion of PduQ is not lethal when grown solely on 1,2-PD, although there is a growth defect. This suggests that internal NAD+/NADH recycling is not strictly necessary for MCP function but may be useful for maintaining NAD+ concentrations in the MCP.

Since PduQ is not required for growth on 1,2-PD, we hypothesized that NADH is able to cross the compartment barrier. This, in turn, implies that internal recycling is not strictly required for MCP function. To test this idea, we created strains lacking PduP (ΔPduP), PduQ (ΔPduQ) or both PduP and PduQ (ΔPduPQ) on the genome. However, work from Bobik and coworkers suggests that PduQ may interact with and be co-encapsulated with PduP^16^. This implies that PduQ encapsulation may be prevented in a PduP knockout, complicating any results from our assay. To investigate if PduQ is still encapsulated in the PduP knockout strain, we developed a FLAG and GFP-tagged PduQ reporter. Using this reporter, we determined that PduQ is still encapsulated when PduP is not present (Figure SI4). We therefore evaluated purified WT, ΔPduP, ΔPduQ, and ΔPduPQ MCPs in our *in vitro* assays to see if the MCPs functioned metabolically without the ability to internally recycle NAD+/NADH (Figure 3). We hypothesized that if PduP or PduQ were not present in the MCP, recycling could not occur. Instead, the production of downstream products would be limited by the amount of corresponding cofactor added.

**Figure 3:**
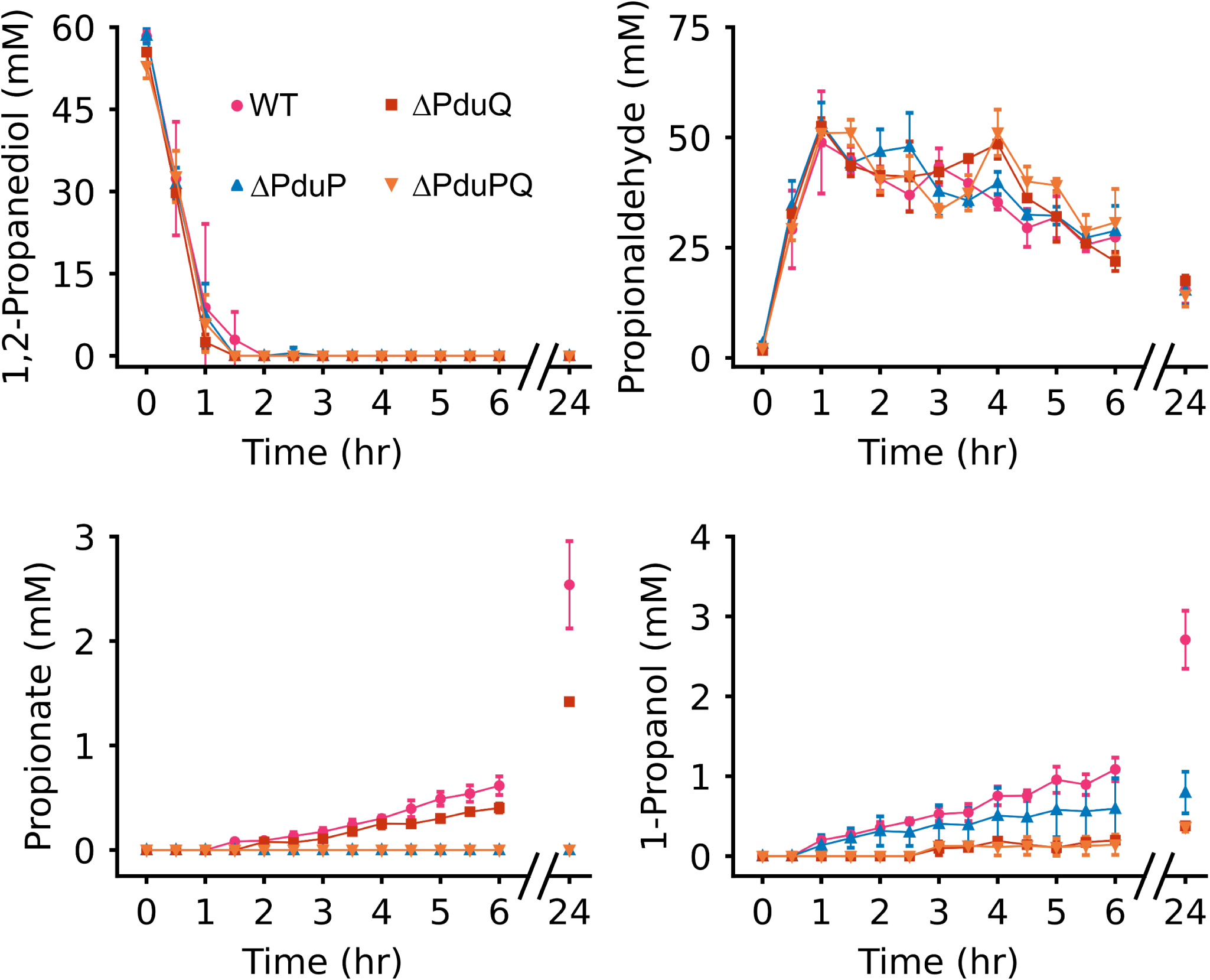
NAD/NADH cofactor recycling is beneficial but not necessary for Pdu pathway flux. In vitro measurement of metabolites corresponding to wild type (magenta), ΔPduP (blue), ΔPduQ (red) or ΔPduPQ (orange) MCPs. Error bars indicate standard deviation across three biological replicates.

Across all the MCP conditions, the 1,2-PD consumption and propionaldehyde production did not vary. This is expected as the first step of the Pdu pathway is not NAD+/NADH cofactor dependent. The apparent similar consumption of propionaldehyde between conditions is surprising given the removal of the downstream enzymes. This may be due to the majority of propionaldehyde being lost to evaporation (Figure SI5). The ΔPduP MCPs did not produce any propionate, and ΔPduQ MCPs had a large reduction in 1-propanol production, which is expected as the enzymes to catalyze the corresponding reaction are not present (Figure 3). The 1-propanol present in the ΔPduQ MCP condition could be due to spontaneous conversion of propionaldehyde to 1-propanol. Interestingly, the ΔPduP MCPs were able to produce detectable 1-propanol but produced less 1-propanol than the WT MCP throughout the entire time series. The rate (mM/hr) of 1-propanol production was higher in the WT MCP condition than in the ΔPduP MCP condition from 0-6 hours (*p-value* = 0.1568) (SI Table 2). In addition, the amount of 1-propanol produced at 24 hours was significantly higher (*p-value* = 0.0080) in the WT MCP than in the ΔPduP MCP condition. Similarly, the ΔPduQ MCPs are able to produce detectable propionate at 1.5 hours which is 30 minutes slower than the WT MCPs, and the amount of propionate produced was also less throughout the entire time series. Analogous to the 1-propanol results, the amount of propionate produced at 24 hours was significantly lower in the ΔPduQ MCP condition compared to the WT MCP condition (*p-value* = 0.0025). In addition, the rate of propionate production from 0-6 hours was significantly lower (*p-value* = 0.0415) in the ΔPduQ MCPs than in the WT MCP condition (SI Table 2). As expected, our negative control ΔPduPQ MCPs produced propionaldehyde, little 1-propanol similar to the ΔPduQ MCP condition, and no propionate. We conclude that while NAD+/NADH cofactor recycling is not necessary for Pdu pathway production of 1-propanol and propionate *in vitro*, it does enable higher production rates and final titers of these products.

### Cofactor recycling helps maintain propionate production in cofactor imbalanced environments

Lower propionate and 1-propanol production in reactions with NAD+/NADH cofactor recycling knocked out suggests some benefit of having internal recycling, even though the Pdu pathway has access to external NAD+/NADH cofactor pools. The NAD+/NADH ratio *in vivo* can vary^40^, so we hypothesized that NAD+/NADH cofactor recycling helps maintain a constant NAD+ pool for the PduP branch to consistently convert propionaldehyde to propionate. To test this, we took our WT MCPs and tested them in reaction environments with apparent NAD+/NADH imbalances (Figure 4A) to determine how production of propionate or 1-propanol varied as a function of the ratio. We tested reaction conditions in which only NAD+ or only NADH were added, as well as conditions where the NAD+/NADH ratio was either 10:1 or 1:10, to investigate how extreme NAD+/NADH imbalances affect Pdu pathway flux.

**Figure 4:**
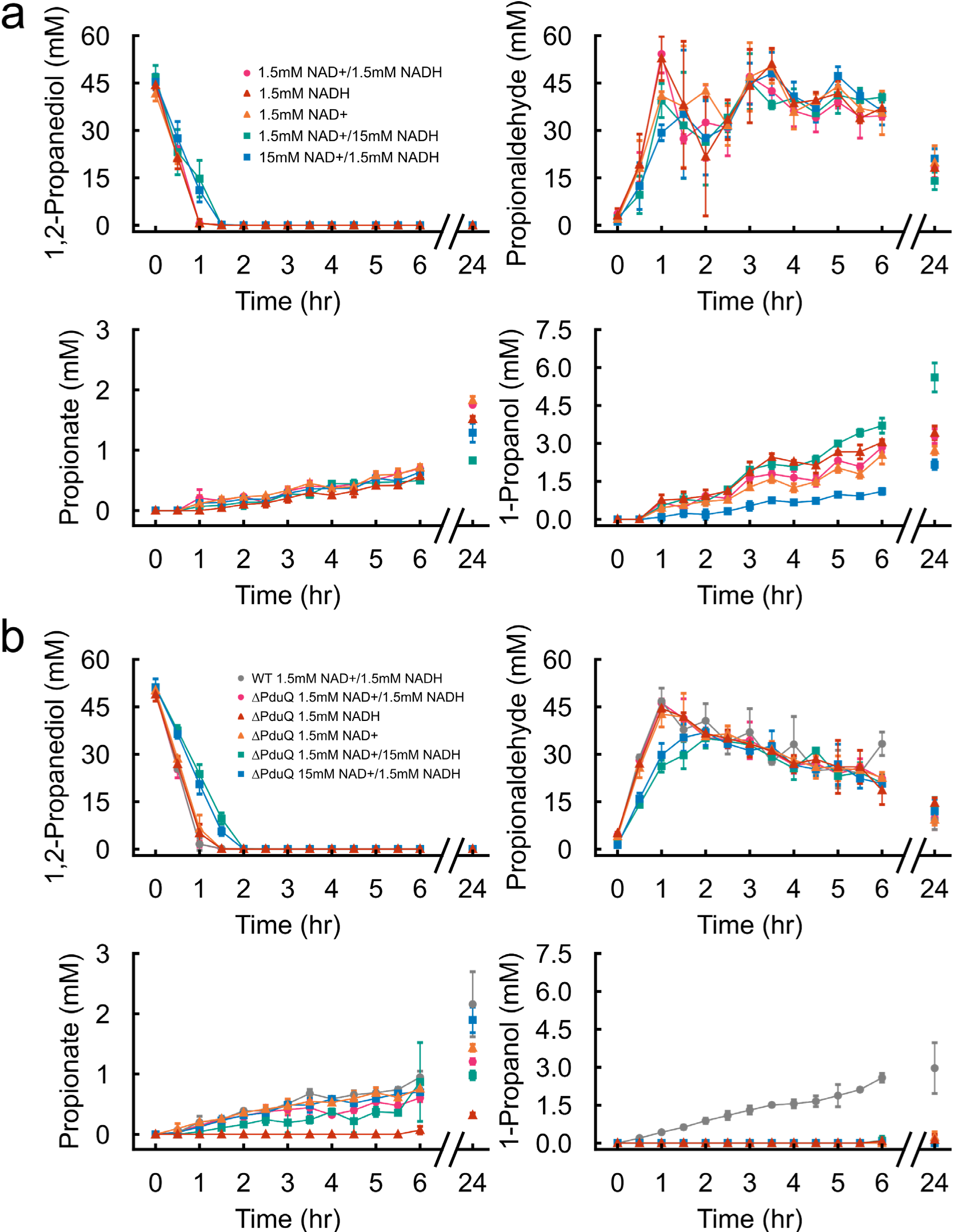
NAD+/NADH cofactor recycling helps maintain propionate production in NAD+/NADH imbalanced environments. **(A)** *In vitro* measurement of metabolites with WT MCPs with 1.5 mM NAD+/NADH, 1.5 mM NAD+ and 15 mM NADH, 15 mM NAD+ and 1.5 mM NADH, 1.5 mM NADH, or 1.5 mM NAD+ added to the reaction. Error bars indicate standard deviation across three technical replicates. **(B)** *In vitro* measurement of metabolites with WT MCPs with 1.5 mM NAD+/NADH, ΔPduQ MCPs with 1.5 mM NAD+/NADH, 1.5 mM NAD+ 15 mM NADH, 15 mM NAD+ 1.5mM NADH, 1.5 mM NADH, or 1.5 mM NAD+ added to the reaction. Error bars indicate standard deviation across three technical replicates.

With NAD+/NADH cofactor recycling present (e.g., with wild-type MCPs present), and when NAD+/NADH was added in a 10:1 or 1:10 ratio, the 1,2-PD consumption was markedly slower than in the other conditions. The rates (mM/hr) of propionate production from 0 – 6 hours were somewhat similar (ANOVA p-value: 0.05880) across the different cofactor environments, but at 24 hours, there was lower propionate production when NAD+ was omitted and even lower when NAD+/NADH ratios were 10:1 and 1:10. This is unexpected given the mechanism of PduP, but may reflect a bottleneck in the pathway downstream of PduP. In contrast, 1-propanol production varies more across the different NAD+/NADH ratios (ANOVA *p-*value = 8.06×10^−9^) than propionate from 0 – 6 hours. As may be expected given that 1-propanol production is directly NADH-dependent, the conditions with only NADH or a 1:10 NAD+/NADH ratio have more 1-propanol production than the conditions with equal amounts of NAD+ and NADH. When NADH is omitted or at a 10:1 NAD+/NADH ratio, 1-propanol production is lower. Collectively, these results suggest that in variable NAD+/NADH cofactor environments, propionate production rates remain largely steady whereas 1-propanol production changes depending on the NAD+/NADH ratio.

The production of propionate was fairly consistent amongst the different NAD+/NADH cofactor environments, so we next sought to investigate to what degree cofactor recycling was helping maintain steady propionate production. To this end, we used ΔPduQ MCPs in the same cofactor balanced or imbalanced environments and collected the metabolite profiles (Figure 4B). We reasoned that the NADH recycling-deficient ΔPduQ MCPs would not be able to regenerate any NAD+ and thus propionate production would be limited by the starting NAD+ amount rather than the total NAD+/NADH pool. Indeed, the rate of propionate production by ΔPduQ MCPs is affected by the relative amount of NAD+ to NADH in the reaction environment from 0 – 6 hours (SI Table 2). In the presence of 1.5 mM NAD+, the rate of propionate production is significantly lower when NADH is present than when only NAD+ is present (p-value = 0.010), and the rate of propionate production decreases as the concentration of NADH is increased (p-value = 0.07339) (SI Table 2). The 1.5 mM NAD+ / 1.5 mM NADH condition does have a higher rate of propionate production than the 1.5 mM NAD+ / 15 mM NADH condition, suggesting lower NADH amounts may result in increased propionate production. The stark difference in propionate production in different NAD+/NADH environments between WT Pdu MCPs and ΔPduQ MCPs suggests that NAD+/NADH cofactor recycling does help keep propionate production consistent in NAD+/NADH imbalanced reaction environments.

### Parameter estimation using systems model supports prior shell permeability predictions

The permeability of the MCP shell to metabolites is fundamental to pathway function, with previous systems modeling predicting permeability–expressed as a diffusive velocity–to be on the order of 10^−5^ cm/s^21^. We calibrated our model to the time course data of metabolites with varying NAD+ to NADH ratios (Figure 4A). This was intended to estimate shell a permeability value that was robust across a range of behaviors under varying cofactor balances.

The experiments succeeded in sampling the desired points in progress space, which were indicated by points of high variability in the experimental design (Figure 2C). Thus we expect the experimental data to be informative for determining k_p_. Time courses for each condition were transformed into progress space, assuming they have reached equilibrium at t=24 hours (Figure 5A). Sampling at the recommended time intervals from t=0.5 hours to t=6 hours corresponded to 40-80% reaction progress to steady-state (grey) for WT and single-cofactor conditions (1.5mM NADH, 1.5mM NAD+), and constrained the permeability near 10.^−4.5^ cm/s with high probability (Figure 5B). While there are a broad range of permeabilities 10^−4^-10^−6^ cm/s that have high probability, values outside this range are overwhelmingly unlikely. Effects of the remaining model parameters are marginalized out, isolating the effect of permeability (k_p_).

**Figure 5:**
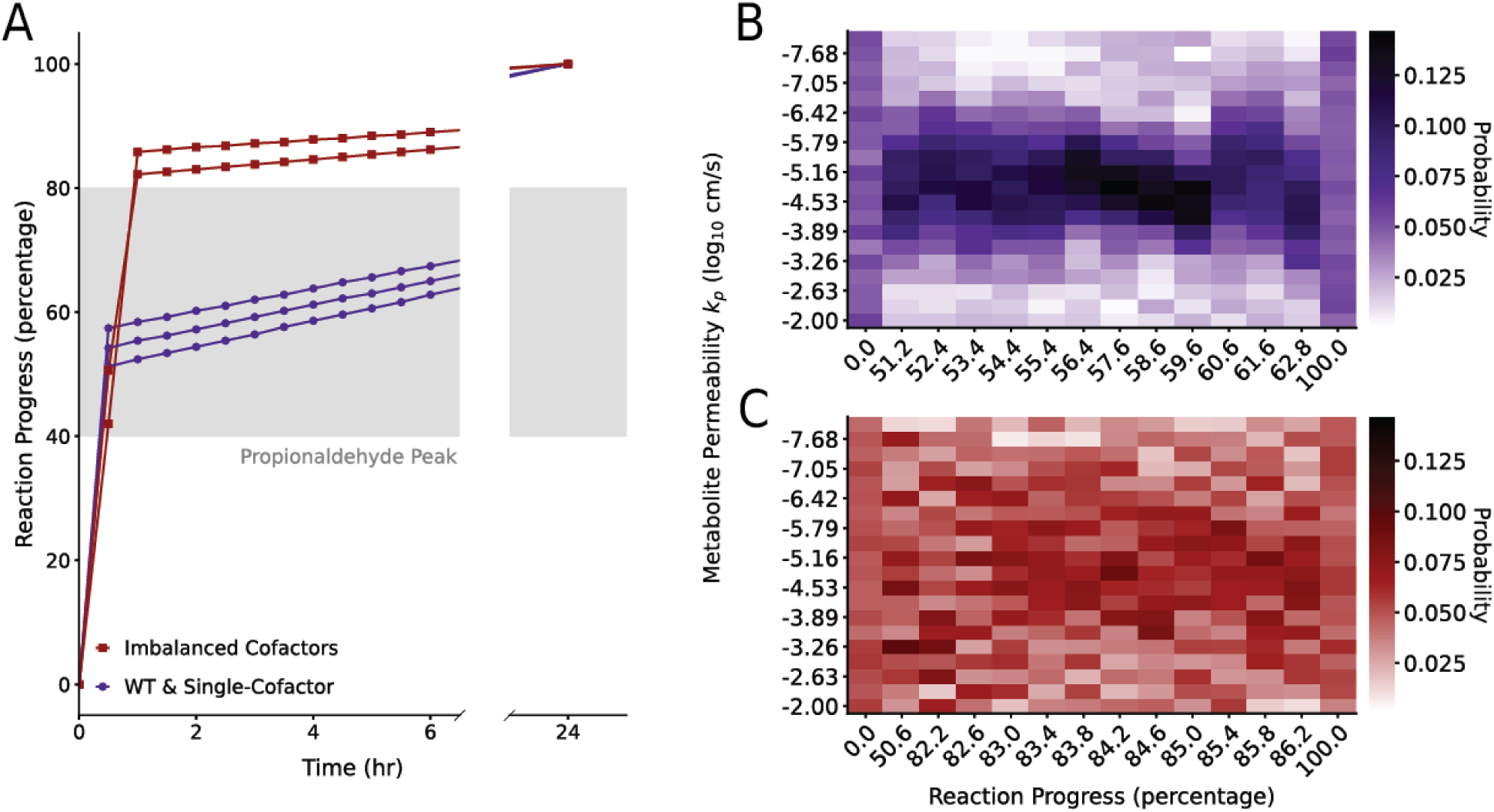
Experimental measurements adhere to model-guided sampling for WT and single-cofactor conditions, constraining estimates of permeability, kp. **(A)** Transformation of time to reaction progress coordinates confirms that the desired progress regime (associated with the propionaldehyde peak) is successfully sampled for WT and single-cofactor conditions (purple) but not for imbalanced cofactor conditions (red). **(B)** Constraint of permeability kp in WT conditions as a progress-dependent probability distribution shows strong constraint to values in the range of 10-6-10-4 cm/s. **(C)** Metabolite permeability kp is poorly constrained in 10:1 NAD+/NADH conditions.

In contrast, in experiments with 10:1 and 1:10 NAD+/NADH ratio conditions we did not match the designed sampling of progress space and only a single sample (t=0.5 hours) fell within 40-80% reaction progress. All samples beyond t=0.5 correcitsponded to the final 20% of reaction progress and the dynamics were not well resolved. The discrepancy between desired and actual sampling in progress space arose because we erroneously assumed that the overall timescale of the dynamics would remain the same across experimental conditions. During all simulations used to inform sampling (Figure 2), NAD+/NADH were held at the WT concentrations (1.5mM). However, large NAD+/NADH imbalances strongly effect the dynamics, changing the timing of the peak and invalidating the experimental design for these conditions. Data collected from these conditions do not constrain estimates of shell permeability in model calibration with many high and low probability values spread over several orders of magnitude (Figure 5B). Further, trends across progress show that even in WT and single-cofactor conditions, where the experimental design is valid, measurements at certain times are uninformative for constraining estimation of permeability (Figure 5C). For example, at both 0% and 100% reaction progress to steady-state, the probability of permeability across its tested range is relatively uniform. Such uniformity provides little information that can refine estimates of permeability from its originally tested range. In all, this shows naive sampling which did not resolve important dynamical features could not inform the permeability, but when sampling followed the experimental design and resolved propionaldehyde peak it was informative and successfully constrained the permeability value.

While experimental design succeeded, simulations from the model with the best-fit parameters recapitulate the upstream 1,2-propanediol and propionaldehyde dynamics but fail to capture the downstream 1-propanol and propionate dynamics (Figure 6A). We hypothesize that the model does not predict downstream dynamics because the model is missing mechanisms to capture the possible inhibition of PduL/P/Q enzymatic reactions (see Discussion). Notably, this did not impact our ability to estimate the permeability. While propionaldehyde concentrations are near the peak, measurements of the upstream metabolites dominate variation (Figure 2) and therefore, the upstream metabolites are critical to informing permeability and downstream dynamics are much less informative. Across 30 initial seeds, the best-fit parameters were found by minimizing the squared error summed across the pooled NAD+/H conditions for propionate and propionaldehyde only. We used a covariance matrix adaptation evolution strategy with an incremental population increase restarts and learning rate adaptation (LRA IPOP-CMA-ES in Python), which is suitable for this nonlinear, high-dimensional system^39–41^. Using only the upstream metabolites in fitting produced comparable results to using all measured metabolite data (Figure 6B). While inclusion of the downstream metabolites in calibration appeared to improve fit quality of the best-fit parameter set, the resultant value of permeability remains fundamentally static. The estimated parameter values are unchanged (diagonal) regardless whether we include the downstream metabolite data in calibration (Figure 6C). Model parameters describing upstream kinetics fall closely to the diagonal while parameters associated with downstream kinetics have more variation. Downstream kinetics, however, are not critical to estimation of permeability. Together, this analysis provides confidence in the resulting estimate of MCP shell permeability at 10.^−4.85^ cm/s found to be optimal by a fit to upstream data. This value, the generalized diffusion velocity of metabolites through the shell, is consistent with estimates that optimize permeability for trapping of metabolites *in vivo* such that 1,2-PD uptake is optimized, while non-toxic levels of propionaldehyde are controlled in the cytosol^21^.

**Figure 6:**
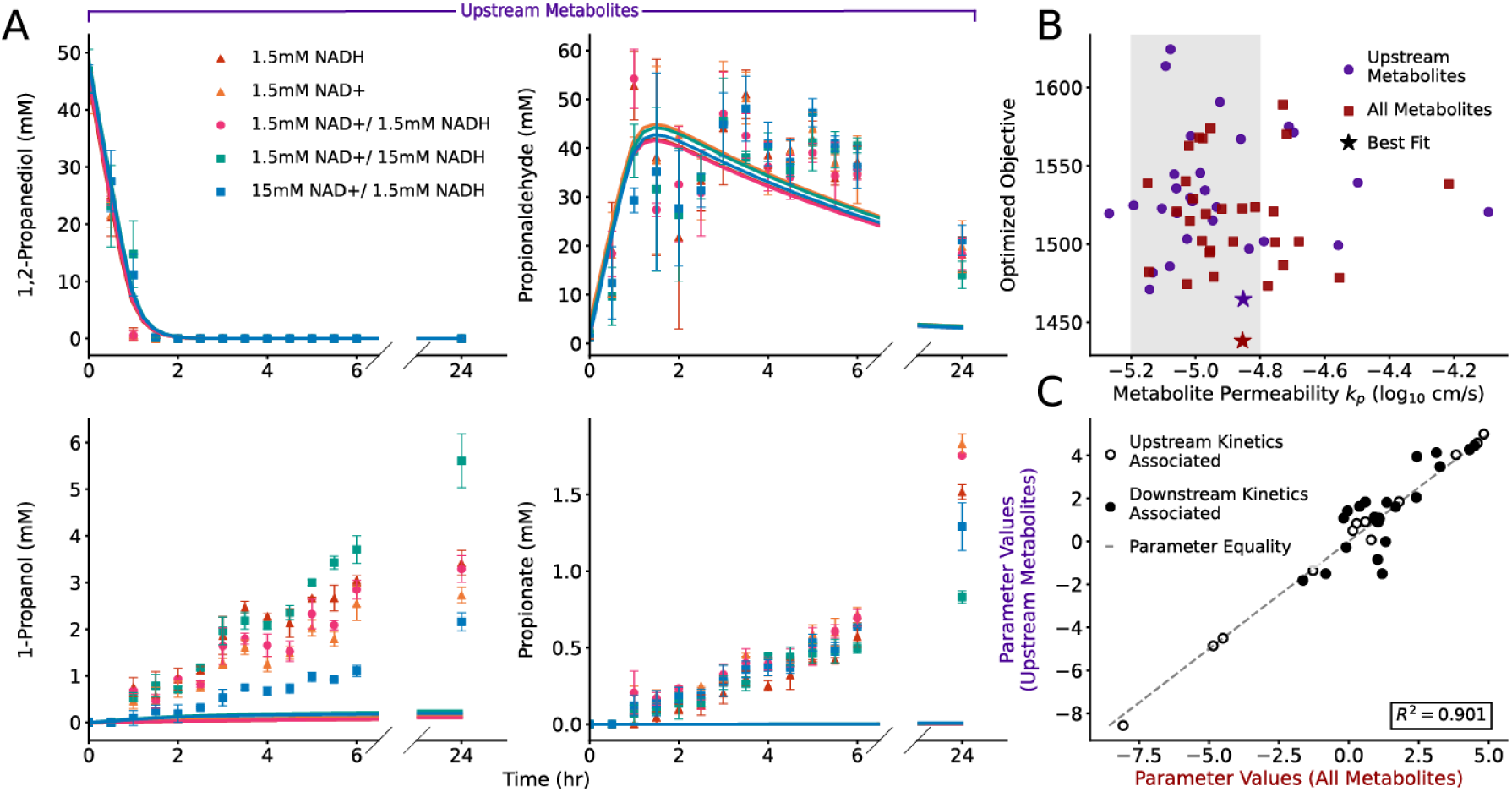
Fit of kinetic model to data captures upstream dynamics. Estimates of permeability and other parameters unimpacted by downstream fit quality. **(A)** Forward simulation with best-fit kinetic parameters fits upstream metabolites (important to capture as per model-guided sampling) but fails to capture end products. **(B)** Parameter estimation using upstream metabolite data (purple) results in same metabolite permeability k_p as calibration using all metabolite data (red). Grey denotes theoretically reasonable range as per^21^. **(C)** Parameters estimated between conditions in Panel B remain largely equal. Divergence primarily occurs in parameters associated with downstream kinetics (i.e., PduP, PduQ, PduL, and PduW/AckA) (closed circles).

## Discussion

In this study, we developed a new *in vitro* assay to investigate the pathway performance of the Pdu MCP. The optimal timepoints for this assay were determined using sampling guided by a systems model predictions of the metabolites and time points with large sensitivity (variation) to changes in unknown parameters. With this assay, we measure multiple metabolites of the Pdu pathway simultaneously and investigate manipulations to both the Pdu MCP enzymes and to the reaction environment. Because the open reaction environment allows for easy manipulation of the system, we can evaluate reaction conditions that are inaccessible by *in vivo* techniques. We used this assay to demonstrate that NAD+/NADH cofactor recycling is beneficial in increasing pathway performance and keeping propionate production consistent. In addition, this assay was used to inform kinetic models, allowing us to estimate properties such as Pdu MCP shell permeability, which is difficult to estimate with experimental work alone.

While there is full pathway activation in this *in vitro* assay, the mass balance does not close as expected. This may be because we were unable to measure several intermediates, including propionyl-CoA and propionyl-PO_4_, and/or due to the loss of some propionaldehyde to evaporation over the course of the reaction (Figure SI5). Notably, the amount of propionaldehyde produced is significantly higher than what has been reported *in vivo*^16,23,27,28,42–45^. More investigation is required to determine why there is such a discrepancy between *in vivo* and *in vitro* metabolite profiles. It is possible that cytosolic enzymes, such as promiscuous alcohol dehydrogenases, aid in processing propionaldehyde, resulting in lower concentrations *in vivo*^18^.

Using this assay, we were able to investigate NAD+/NADH recycling within compartments. Our data suggests that while NAD+/NADH recycling is beneficial to Pdu pathway activity, it is not required for either propionate or propanol production in an *in vitro* environment. This suggests that the benefits of NAD+/NADH cofactor recycling outweigh the possible benefits of diverting all of the Pdu pathway flux to propionate. If this behavior also applies *in vivo,* it could help explain the growth deficit observed when NAD+/NADH cofactor recycling is removed.

The WT Pdu MCPs produced similar amounts of propionate amongst the different NAD+/NADH cofactor environments. However, when using ΔPduQ MCPs (no NAD+ recycling) in the same cofactor imbalanced environments, propionate production was much lower in limited NAD+ environments. The presence of NADH also reduced propionate production when using ΔPduQ MCPs (no NAD+ recycling). Taken together, our results suggest that cofactor recycling plays a role in maintaining an appropriate NAD+ concentration or NAD+/NADH ratio for optimal propionate production in this *in vitro* environment. Notably, PduQ both consumes NADH and produces NAD+, each of which pushes the PduP reaction in the “forward” direction toward propionyl-CoA production. Keeping propionate production constant is evolutionarily beneficial since this would result in a consistent carbon source. It is worth noting that we do not measure propionyl-CoA or propionyl-PO_4_, but the overall production of propionate may be influenced by the downstream flux through the branch, in addition to the NAD+ pool for PduP.

The 1:10 and 10:1 ratios of NAD+/NADH were higher than what has been recorded in *E. coli*^46^. In these environments, WT Pdu MCPs were still able to produce similar propionate from hours 0-6, suggesting the Pdu MCP pathway can tolerate high NAD+/NADH imbalances. We also find it interesting that in the 1:10 and 10:1 NAD+/NADH environments, 1,2-PD consumption and propionaldehyde production were slower. Why this step slows down in an environment with higher available cofactor concentration is a phenomenon that requires further investigation.

We used this assay in combination with mathematical modeling to estimate the shell permeability to all metabolites, a key physical parameter that has been predicted in previous modeling work but has yet to be estimated from data. As the permeability of MCPs is likely selective, this represents an order of magnitude estimate, and further work would be needed to determine differential permeabilities for metabolites in the Pdu pathway^30^. By assuming an *in vitro* setting, we simplified our model structure and calibration process in comparison to *in vivo* settings. Our simple model was able to reasonably capture the upstream metabolite dynamics needed to estimate shell permeability, but the model was unable to recapitulate the low production of downstream metabolites observed in experiments. Despite exploring several methodologies, including overweighting the final time points and normalizing the data to more heavily weight metabolites with lower concentrations, we could not find a consistent model fit for high 1,2-PD consumption and relatively low conversion to the downstream metabolites. Neither reversibility of the enzymes nor sequestration in the unmeasured intermediate propionyl-CoA, both of which are captured in the model, were sufficient. This points to an unmodeled mechanism that slows the downstream catalysis of propionaldehyde by the PduQ and/or PduP-PduL-PduW/AckA branches. Potential mechanisms for this phenomena include enzyme denaturation in either branch, oxidation of PduQ, or CoA inhibition of^17,46–49^. Further investigation is needed to determine if the inclusion of enzyme sequestration effects in the model allows for closer alignment to observed data.

With this *in vitro* Pdu MCP system combined with kinetic modeling, we have been able to probe properties of the Pdu MCP, such as Pdu MCP shell permeability, not accessible with *in vivo* studies. This system can also be used as a testbed for different enzymatic configurations within the Pdu MCP or even the performance of encapsulated heterologous pathways. Overall, we have developed a new tool that can be used to more directly study the Pdu MCP in isolation as well as get closer to determining what properties of a pathway will make it more likely to benefit from encapsulation.

## Methods

### Bacterial strains and plasmids

*Salmonella enterica* serovar Typhimurium LT2 was used for the generation of 1,2-propanediol utilization MCPs (Pdu MCPs) gene knockouts, purification of Pdu MCPs, and assessment of PduQ encapsulation in the Pdu MCP via fluorescent reporters. All of the strains used in this study can be found in Table SI1A with details on their genotype and any plasmids they harbor. All primers used in this study can be found in Table SI1B.

### Strain generation (recombineering)

Genomic modifications to strains were made using the λ red recombineering technique as previously described^50^. Briefly, LT2 was transformed with the plasmid pSIM6 which encodes the λ red recombineering machinery and carbenicillin resistance. The pSIM6 plasmid is ejected from the cell at 37 °C and expresses the λ red recombineering genes at 42 °C. Electroporation was used to introduce a cassette containing a chloramphenicol resistance gene (*cat*) and a sucrose sensitivity gene (*sacB*) amplified from the TUC01 genome with added, sequence-specific homology to a target locus. Cassette incorporation at the locus of interest was confirmed by growing cells on lysogeny broth (LB)-Agar supplemented with 10 μg/mL of chloramphenicol and 30 μg/mL of carbenicillin at 30 °C. Single colonies from the chloramphenicol and carbenicillin plates were streaked onto 6% (w/w) sucrose plates to determine if the strains were sucrose sensitive. The *catsacB* cassette was replaced with either single stranded DNA for knockouts or a PCR product of a full gene with homology to the locus of interest. Incorporation of the DNA of the desired gene was screened with sucrose sensitivity, and polymerase chain reaction was conducted at the locus of interest and sent for Sanger sequencing (Genewiz).

### Pdu MCP Purification

Pdu MCPs were purified using a differential centrifugation method as previously described^51,529^. To purify Pdu MCPs, we inoculated 5 mL of LB liquid media with LT2 and grew the cultures at 30 °C, 225 rpm for 24 hours. The overnight culture was subcultured 1:1,000 (v/v) into 200 mL of No Carbon Essential (NCE) media (29 mM potassium phosphate monobasic, 34 mM potassium phosphate dibasic, 17 mM sodium ammonium hydrogen phosphate) supplemented with 50 µM ferric citrate, 1 mM magnesium sulfate, 42 mM succinate as a carbon source, and 55 mM 1,2-propanediol as the *pdu* operon inducer. This culture was grown at 37 °C and 225 rpm until the OD_600_ reached 1-1.5. The cells were then collected via centrifugation and resuspended in a lysis buffer (32 mM Tris-HCl, 200 mM potassium chloride, 5 mM magnesium chloride, 0.6% (v/v) 1,2-propanediol, 0.6% (w/w) octylthioglucoside, 5 mM β-mercaptoethanol, 0.8 mg/mL lysozyme (Thermo Fisher Scientific), 0.04 units/mL DNase I (New England Biolabs, Inc.) pH 7.5–8.0). The resuspended cells were rocked at 60 rpm at room temperature. After 30 minutes, the lysed cells were kept on ice for 5 minutes to stop lysis. Centrifugation at 12,000 xg for 5 minutes was then used to clarify the lysate. The MCPs are anticipated to be in the supernatant fraction of the clarification spin, which is then taken and subjected to additional centrifugation at 21,000 xg for 20 minutes to separate the MCPs from other soluble cellular proteins. Following this spin, the pellet will be enriched for MCPs so the supernatant was removed, and the pellet was washed with buffer (32 mM Tris-HCl, 200 mM KCl, 5 mM MgCl2, 0.6% (v/v) 1,2-propanediol, 0.6% (w/w) OTG, pH 7.5–8.0). MCPs were pelleted again and then resuspended in buffer (50 mM Tris-HCl, 50 mM KCl, 5 mM MgCl2, pH 8.0) and stored at 4 °C until use. A bicinchoninic acid assay was used to measure the concentration of the purified Pdu MCPs (Thermo Scientific).

### Transmission electron microscopy

For transmission electron microscopy, a glow discharge system was used to hydrophilize 400-mesh copper grids with Formvar/carbon film (EMS Cat# FCF400-Cu-50). A 10 µL sample of the purified Pdu MCP sample was then placed onto the grid. Sample was wicked away after 5–10 seconds, and then 10 µL of a 1% (w/v) aqueous uranyl acetate (UA) solution was added as a negative stain. The UA was immediately wicked away. This process was repeated twice for additional UA applications. The first was wicked away immediately, and the second was allowed to sit with the UA for 4 minutes before wicking. A JEOL 1400 Flash transmission electron microscope equipped with a Gatan OneView Camera was used to image the grids at 50,000 x magnification at room temperature (Figure SI5). ImageJ was used to crop the images.

### Phase contrast and fluorescence Microscopy

Fluorescence patterning of the GFP-PduQ reporter was analyzed using phase contrast and fluorescent microscopy. Cells were grown from a single colony overnight in LB-media at 37 °C, 225 rpm for 16-20 hours. This overnight culture was then subcultured 1:100 into NCE media supplemented with 50 µM ferric citrate, 1 mM magnesium sulfate, 42 mM succinate as a carbon source, and 55 mM 1,2-propanediol as the *pdu* operon inducer. The subculture was allowed to grow at 37 °C, 225 rpm 18 hours. 1 mL of subculture was spun down at 4,000 xg for 90 seconds, and 800 µL of supernatant was removed. The cell pellet was resuspended in the remaining 200 µL of supernatant. 1.48 µL of cells were pipetted on Fisherbrand™ frosted microscope slides (Thermo Fisher Scientific Cat# 12-550-343), and sandwiched between the slide and a 22 × 22 mm, #1.5 thickness coverslip (VWR Cat# 16004-302). Both the microscopy slides and slide covers were cleaned with 70% ethanol prior to use. The cells were then imaged on a Nikon Eclipse Ni-U upright microscopy using a 100X oil immersion objective using an Andor Clara digital camera. Image acquisition was done on the NIS Elements Software (Nikon). A C-FL Endow GFP HYQ bandpass filter was used to acquire GFP fluorescence. For GFP fluorescence, a 500 ms exposure time was used.

### Fluorescence measurement

To measure GFP fluorescence as a proxy for expression of the GFP-PduQ reporter, cells were grown from a single colony overnight in LB-media at 37 °C, 225 rpm for 16-20 hours. This overnight culture was then subcultured to an OD600 of 0.05 into NCE media supplemented with 50 µM ferric citrate, 1 mM magnesium sulfate, 42 mM succinate as a carbon source, and 55 mM 1,2-propanediol as the *pdu* operon inducer. This subculture was grown at 37 °C, 225 rpm for 24 hours. 200 µL of culture was pipetted into a 96-well black flat bottom plate (Costar) and ran on a Synergy H1 plate reader (Biotek). GFP fluroescence was excited at 485 nm and emission was measured at 516 nm.

### Coomassie and western blotting

Pdu MCPs were run on sodium dodecyl sulfate-polyacrylamide gel electrophoresis (SDS-PAGE) and then analyzed with Coomassie-blue staining or anti-FLAG western blotting to determine the contents of the Pdu MCP. Purified Pdu MCP samples were diluted to a concentration of 150 ug/mL in buffer (50 mM Tris-HCl, 50 mM KCl, 5 mM MgCl2, pH 8.0). 4 x Laemmli buffer with 10% β-mercaptoethanol was added to a final concentration of 112.5 ug/mL Pdu MCPs, 1 x Laemmli buffer, and 2.5% β-mercaptoethanol. The diluted Pdu MCPs were boiled at 95 °C for 15 minutes. Samples were loaded onto a 15% SDS-PAGE gel (1 ug for Coomassie, 1.5 ug for western blotting). SDS-PAGE gels for Coomassie-blue staining were ran at 120 V for 100 minutes. These gels were then stained with Coomassie Brilliant Blue R-250 and then imaged on an Azure 600 imaging system.

To perform western blots, SDS-PAGE gels were loaded with sample and run at 150 V for 1 hour. To transfer samples to a polyvinylidene fluoride (PVDF) membrane made specifically for fluorescent secondary antibodies (Immobilon), a Bio-Rad Transblot SD was used and set at 25 V, 150 mA, for 35 min. The buffer TBS-T (20 mM Tris, 150 mM sodium chloride (NaCl), 0.05% (v/v) Tween 20, pH 7.5) with 5% bovine serum albumin (w/w) was used to wash the membranes for 1 hour at room temperature. The membrane was then washed in mouse anti-FLAG M2 antibody (Sigma Aldrich) diluted 1:6,666 in 1% (w/w) bovine serum albumin overnight at 4 °C. The membrane was washed in TBS-T, and then goat anti-mouse IgG polyclonal antibody (IRDye® 680RD) (Licor) was applied to the membrane and incubated at room temperature. Incubation time depended on what secondary antibody was used. For goat anti-mouse IgG (H+L) conjugated to horseradish peroxidase, the incubation was 35 minutes. An Azure 600 using a 680 excitation and 694 emission for fluorescent secondary antibody and chemiluminescent signal was used to detect the goat anti-mouse IgG (H+L) secondary antibody. ImageJ was then used to perform densitometry.

### Cloning AckA

AckA was cloned into an expression vector, expressed in cell culture, and purified using a His affinity column. A sequence encoding AckA with an N-terminal 6 x histidine tag (His-AckA) and flanked by BsaI cut sites was ordered as double stranded DNA (Twist). This sequence was inserted into the pET21b vector using Golden Gate cloning which was confirmed with Sanger sequencing (Genewiz)^50^. AckA was purified with a 6 x histidine tag (His-AckA) as described previously^51^. In short, single colonies were inoculated into LB liquid media supplemented with 50 µg/mL of carbenicillin and grown at 30 °C at 225 rpm for 24 hours. The overnight culture was subcultured 1:100 in 1 L of liquid LB media in 2.5 L flasks and grown at 37 °C at 225 rpm. At an OD600 of 0.4-0.8, 1 mL of 1 M IPTG was added to the cultures to induce His-AckA expression. The subculture was allowed to grow for 2 more hours, and the cells were harvested by centrifugation (13,000 x g, 10 minutes, at 4 °C). The cell pellets were collected into 50 mL conical tubes and frozen at −20 °C overnight. Approximately 5 g of dry cell weight is collected from 1 L of subculture.

### Purification of His-AckA

The His-AckA construct was purified as previously described^51^. Briefly, cell pellets were thawed at room temperature and resuspended in 65 mL of resuspension buffer (50 mM HEPES buffer, 500 mM sodium chloride, 20 mM imidazole, 2 mM magnesium chloride, and 2 mM β-mercaptoethanol, 20 µg/mL DNAse I, 250 µM phenylmethylsulfonyl fluoride, pH 7.4). Resuspended cells were lysed with one pass through an Avestin Emulsiflex C3 homogenizer at 20,000 – 25,000 psi. Lysate was clarified with centrifugation (100,000 x g, 45 minutes, 4 °C). The clarified lysate was then filtered through a 0.45 mM filter. The filtered lysate was loaded onto a His-gravitrap column (Cytiva) equilibrated with 10 column volumes of resuspension buffer. The clarified lysate was passed through the column. The column was then washed 3 times with 10 column volumes of resuspension buffer. The His-AckA was then eluted using 6 column volumes of resuspension buffer with 500 mM imidazole into 6 fractions. Each of the elution fractions were run on an SDS-PAGE gel and protein was visualized using Coomassie stain. The fraction with the most His-AckA was dialyzed using a 3.5 kDa cassette against 4L of dialysis buffer (50 mM HEPES, 150 mM potassium chloride, 2 mM of magnesium chloride, 2 mM DTT). After dialysis, 30 mg/mL sucrose was added and the fraction was lyophilized. Lyophilized His-AckA was stored at −20 °C.

### Testing His-AckA activity

His-AckA activity was evaluated using a modified *in vitro* assay to measure acetyl PO_4_ consumption over time as described previously^51,52^. Briefly, purified His-AckA was diluted 1:100. 20μL of the diluted His-AckA was added to 130μL of prewarmed reaction buffer (100 mM Tris-HCl pH 7.4, 10 mM MgCl_2_, 71 mM ADP, and 50 mM acetyl PO_4_). These *in vitro* reactions were incubated at 37 °C for 0, 2, 4, 6, 8, and 10 minutes. Samples for each timepoint were quenched with 150 μL of 10% TCA. After 10 minutes, 50 μL of 705mM hydroxylamine hydrochloride was added to each reaction, and the reactions were incubated at 60 °C for 5 minutes. After 5 minutes, 100uL of 2.5% Fe(III)Cl_2_ was added to the reactions. The reactions were allowed to incubate for 30 minutes at room temperature. Afterward, 200 μL was loaded onto a clear 96-well plate (Greiner Bio-one) and absorbance was measured at 540nm. Absorbance was compared to a standard curve of acetyl PO_4_. 25 units per mg was generated from the His-AckA purification. Units are defined as how much His-AckA (mg) is required to consume 100μM of acetyl PO_4_ at 37 °C.

### Statistical methods

A two tailed *t*-test (df = 4) was used to investigate any significant differences. Three biological replicates were tested for the WT, ΔPduP, ΔPduQ, and ΔPduPQ MCPs in Figure 3. For figures 4A and 4B, three technical replicates were tested for each cofactor condition.

### Run *in vitro* Pdu MCP assay

*In vitro* Pdu MCP assays were performed in 30 μL reactions in 2 mL Eppendorf tubes and incubated at 30 °C. Figure 1 used purified AckA from Sigma Aldrich and consisted of 100 mM Bis-Tris buffer, acetate salts (8 mM magnesium acetate, 10 mM ammonium acetate, 134 mM potassium acetate), 55 mM 1,2-propanediol, 20 μM adenosyl-B_12_, 1 U AckA (Sigma-Aldrich) and 66.66 μg purified microcompartments. Due to supply chain issues, figures 3 and 4 used His-AckA purified in house consisted of 100 mM Bis-Tris buffer, glutamate salts (8 mM magnesium glutamate, 10 mM ammonium glutamate, 130 mM potassium glutamate), 55 mM 1,2-propanediol, 20 uM adenosyl-B_12_, 4 U/mL His-AckA), 1.5 mM ADP, 1.5 mM K_2_HPO_4_ and 66.66 μg purified microcompartments. 1.5mM ADP and 1.5mM K_2_HPO_4_ were added as the in house purified His-AckA performed similar with ADP and K_2_HPO_4_ to the Sigma AckA (Figure SI6). The purified His-AckA appeared to work longer than the Sigma AckA, however, this did not affect our sampling times. Unless otherwise stated, ATP, CoA, NAD+ and NADH were added at various concentrations. Reactions were quenched at each time point by precipitating proteins using 30 μL of 10% trichloroacetic acid and centrifuging at 21,000 x g for 10 minutes at 4 °C. The resulting supernatant was then stored at −80 °C until analysis by HPLC.

### HPLC

Pdu metabolites were quantified using an Agilent 1260 HPLC system. Samples were stored in skirted 96-well PCR plates (BioRad) with a layer of mineral oil above the sample. The mineral oil prevented evaporation of the propionaldehyde intermediate. 5 μ L of sample was injected and metabolites were separated using a Rezex™ ROA-Organic Acid H+ (8%) LC Column (Phenomenex) at 35 °C with 5 mM sulfuric acid as the mobile phase at a flow rate of 0.4 mL/min for 45 minutes. Metabolites were detected using a refractive index detector (RID) and concentrations were determined using the Agilent ChemLab software by comparing against peak areas of known dilutions of 1,2-propanediol, propionaldehyde, 1-propanol and propionate as described previously^27^. For rate, the slope of metabolites was measured from 0 to 6 hours.

## Acknowledgements

We would like to thank and acknowledge members of the Jewett, Mangan, and Tullman-Ercek group for their helpful discussion and preparation of this manuscript as well as constructive comments during the process of writing this manuscript. We would like to specifically thank Elizabeth Johnson, Nolan Kennedy, Svetlana Ikonomova, Taylor Nichols, Blake Rasor, Andre Archer, and Alasdair Hastewell for their helpful discussions. In addition, we want to thank Elizabeth Johnson for collecting the TEM images used in this study and Reese Richardson for critical figure input.

We also want to thank the NUANCE cores for allowing us to use their facilities for this work.

## Author Contributions

DTE, NMM, MCJ, CHA, and SS conceived this project. DTE, NMM, MCJ, CEM, CHA, and SS provided conceptual and technical advice. BJP, CHA, and CEM performed experiments. CC, SS, and NMM developed the software for a kinetic systems model. All of the authors contributed to the analysis and interpretation of the data. BJP, CC, CHA, SS, DTE, and NMM wrote the manuscript. All of the authors reviewed and contributed to the manuscript.

## Data Availability

Any strains and plasmids used in this article are available upon request. All data used in the metabolite profiles and quantitative analysis are provided in the Source Data file.

## Code Availability

Code will be made available upon publication.

## Supplemental Information

**Supplemental Table 1:**
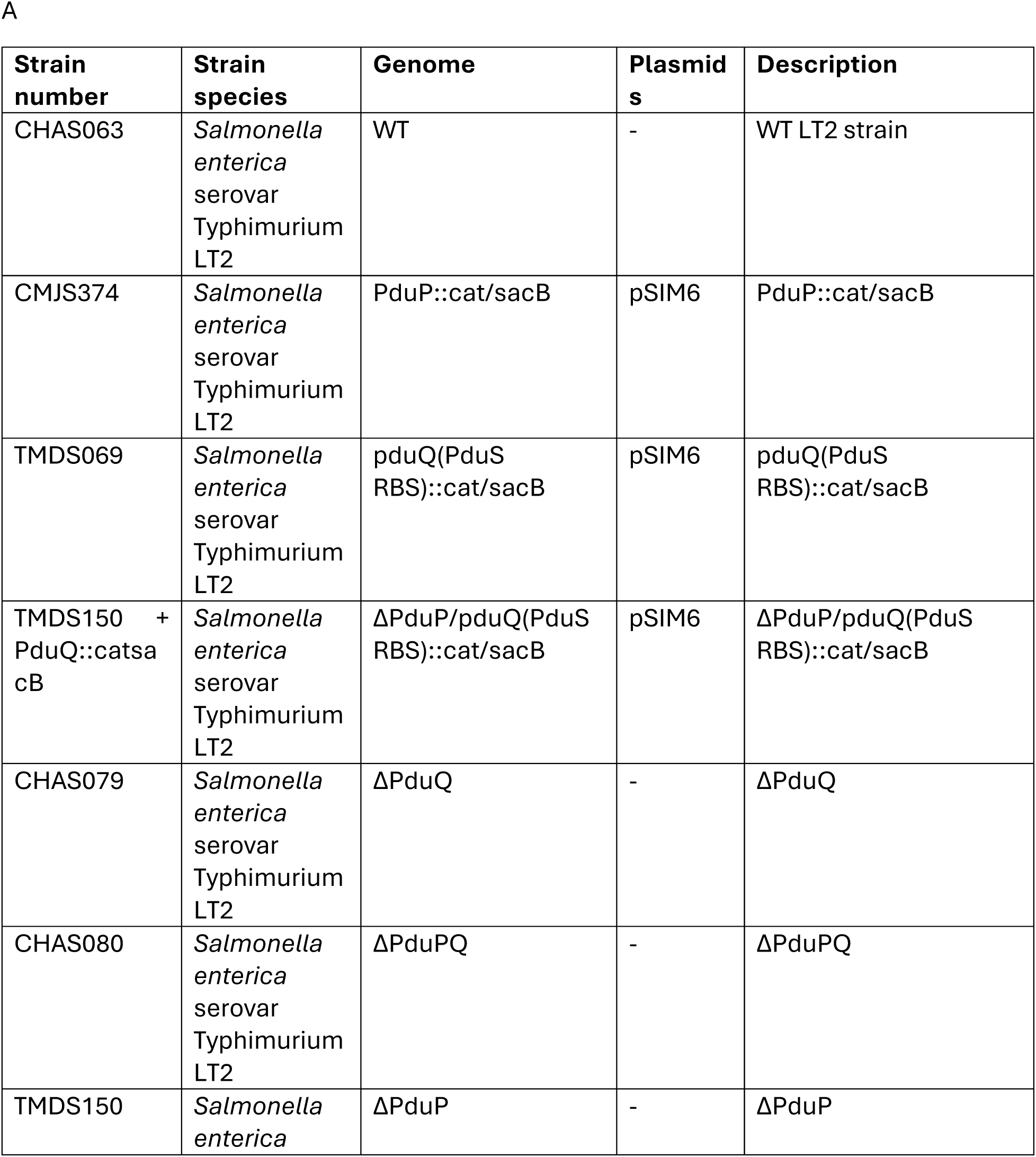
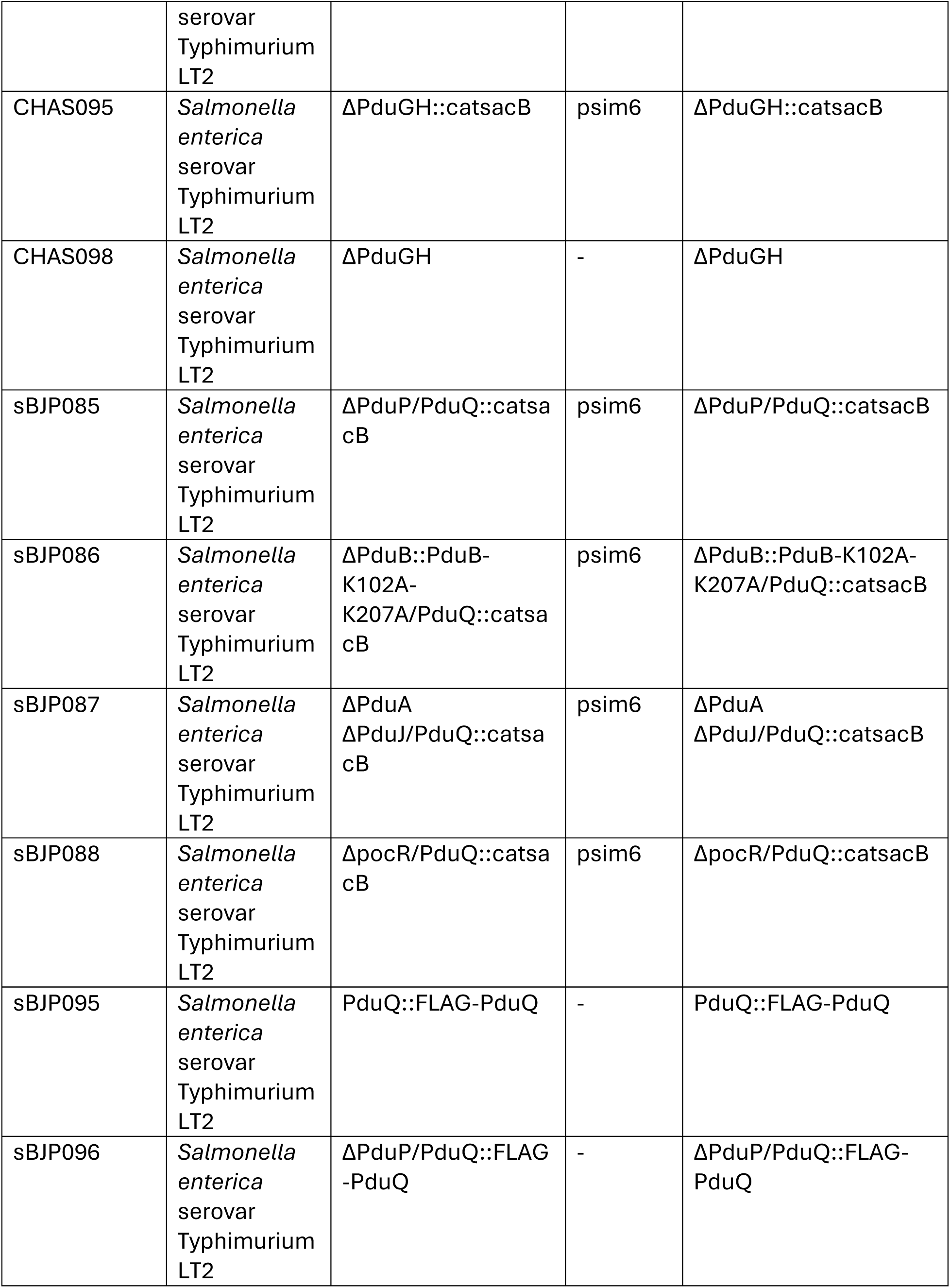
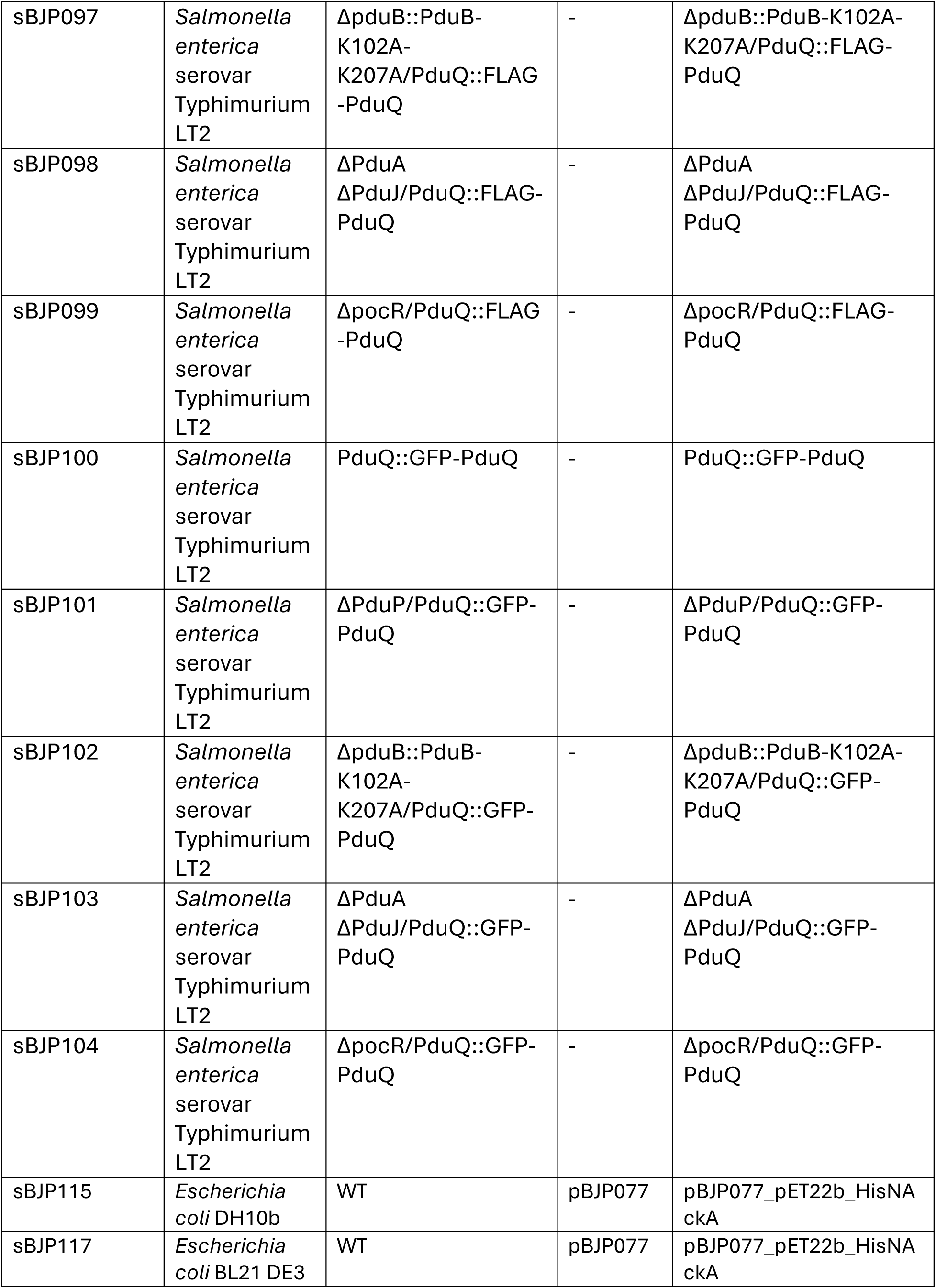
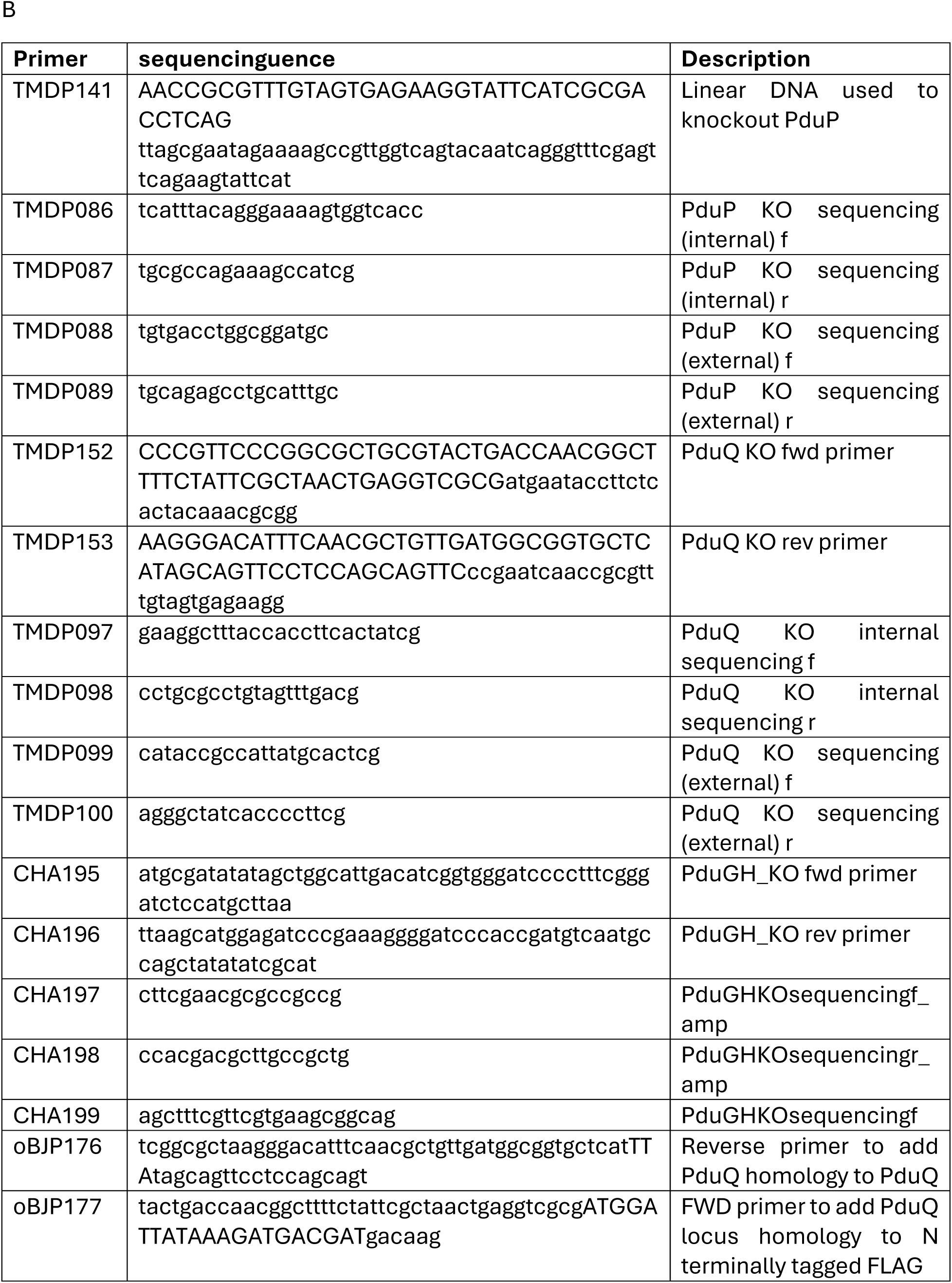
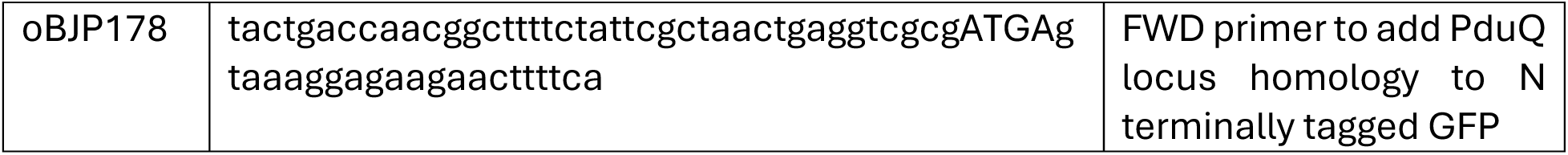
A) Table of strains used in this study. B) Table of primers used in this study.

**Supplemental Figure 1:**
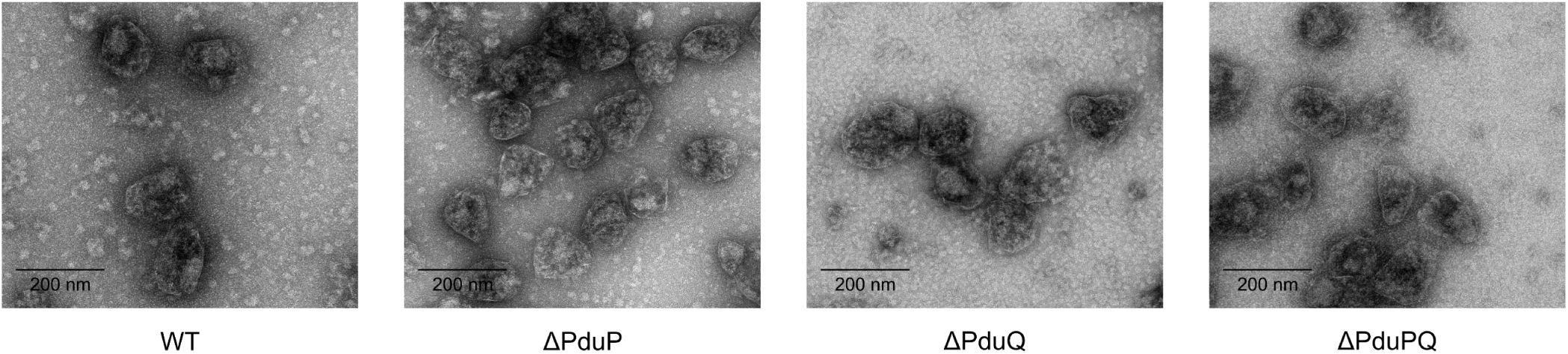
TEM images of purified Pdu MCPs. Transmission electron micrographs of purified WT Pdu MCPs, ΔPduP Pdu MCPs, ΔPduQ Pdu MCPs, and ΔPduPQ Pdu MCPs. Scale bars are 200 nm.

**Supplemental Figure 2:**
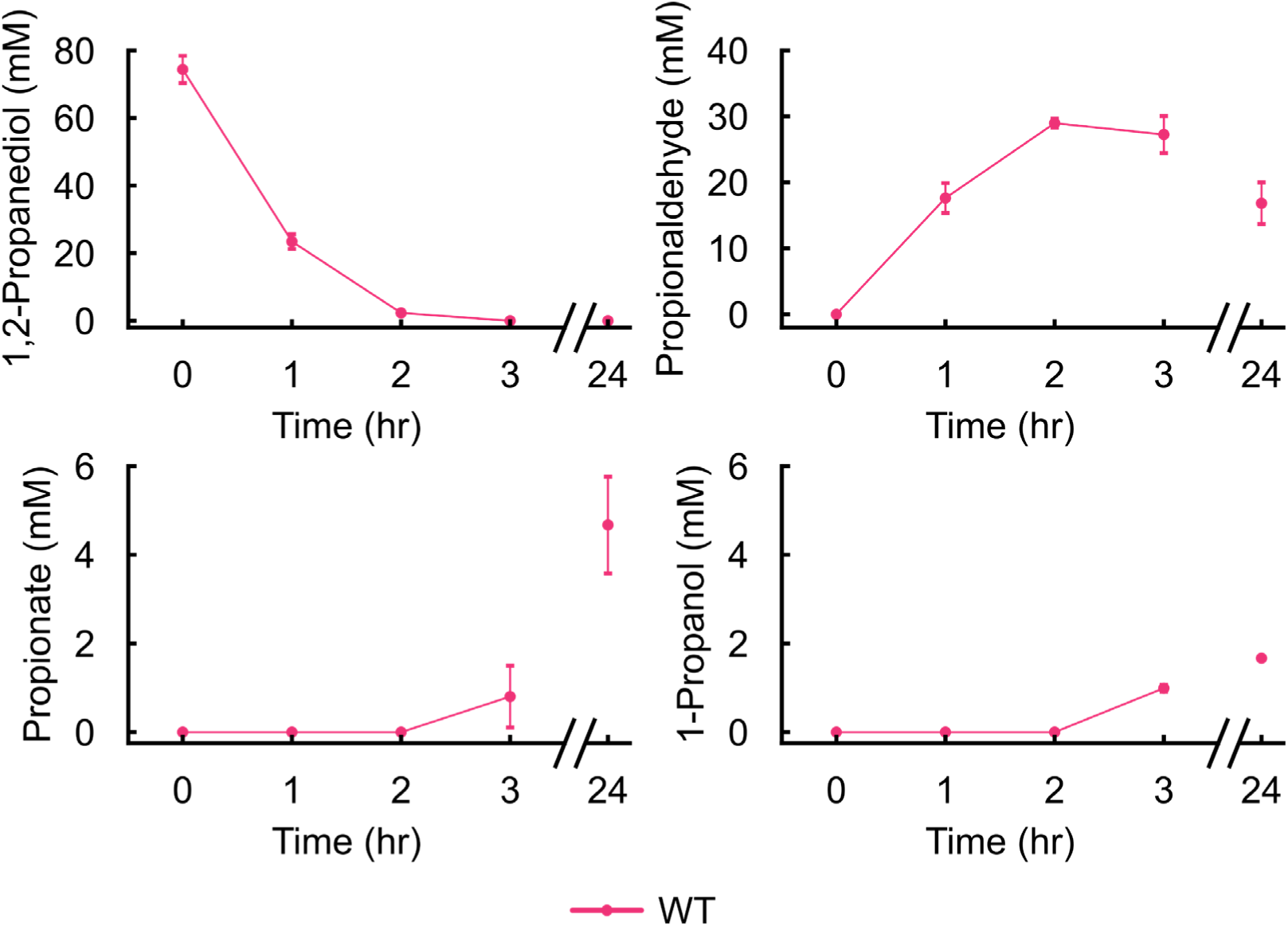
Combining purified Pdu MCPs, 1,2-PD, and the Pdu pathway cofactors results in full pathway activation in vitro. WT Pdu MCPs were added to reaction buffer containing the following: 1,2-PD, B_12_, CoA, NAD+, NADH, and ATP. Concentrations of the metabolites 1,2-PD, propionaldehyde, propionate, and 1-propanol were measured at 0, 1, 2, 3, and 24 hours. Error bars indicate one standard deviation of three technical replicates.

**Supplemental Figure 3:**
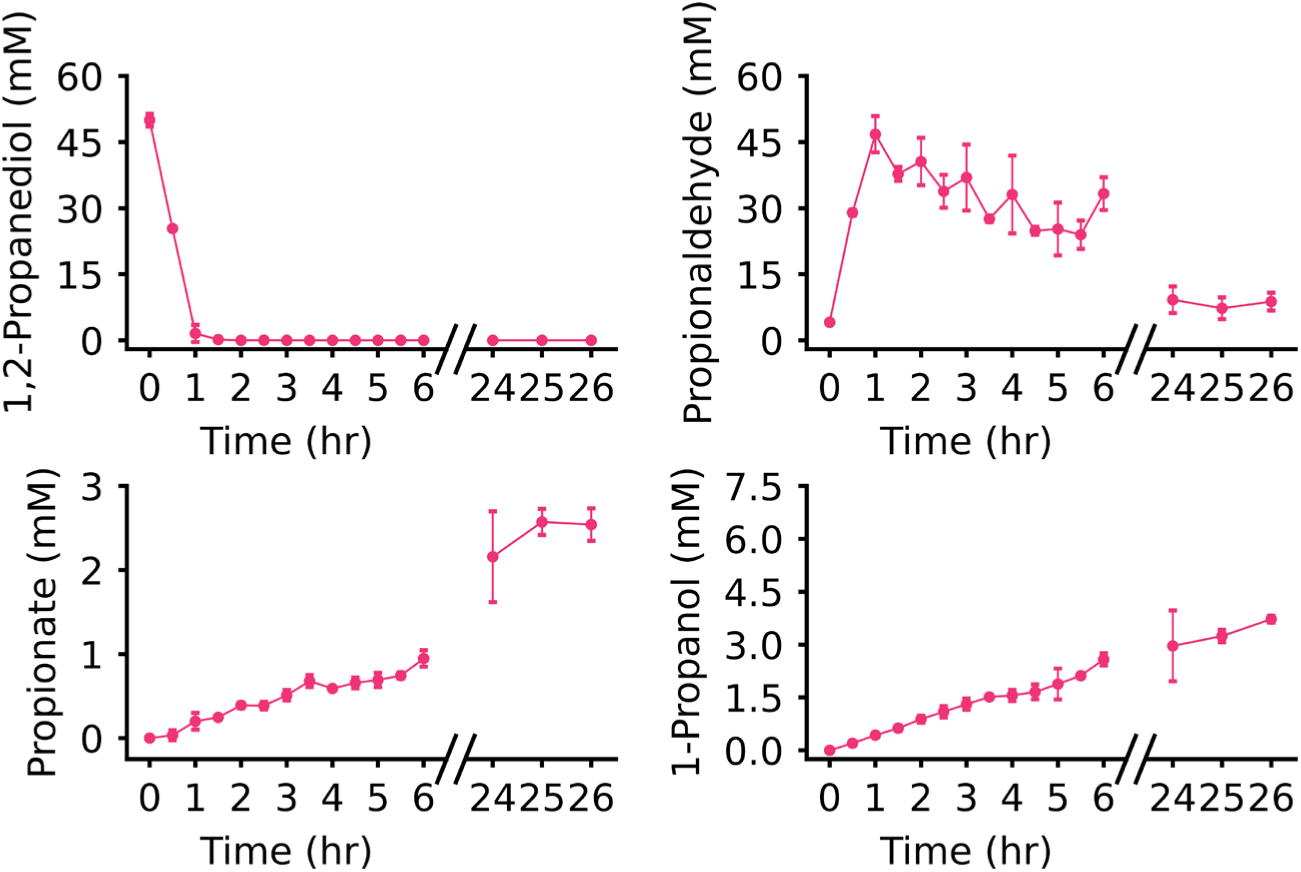
Steady state is reached at 24 hours. WT Pdu MCPs were assayed with measurements taken from 0-6 hours and from 24-26 hours to show steady state at 24 hours. WT Pdu MCP metabolite data from 0-6 hours and 24 hours was also displayed in Figure 4B. Error bars indicated one standard deviation above and below across three technical replicates.

**Supplemental Figure 4:**
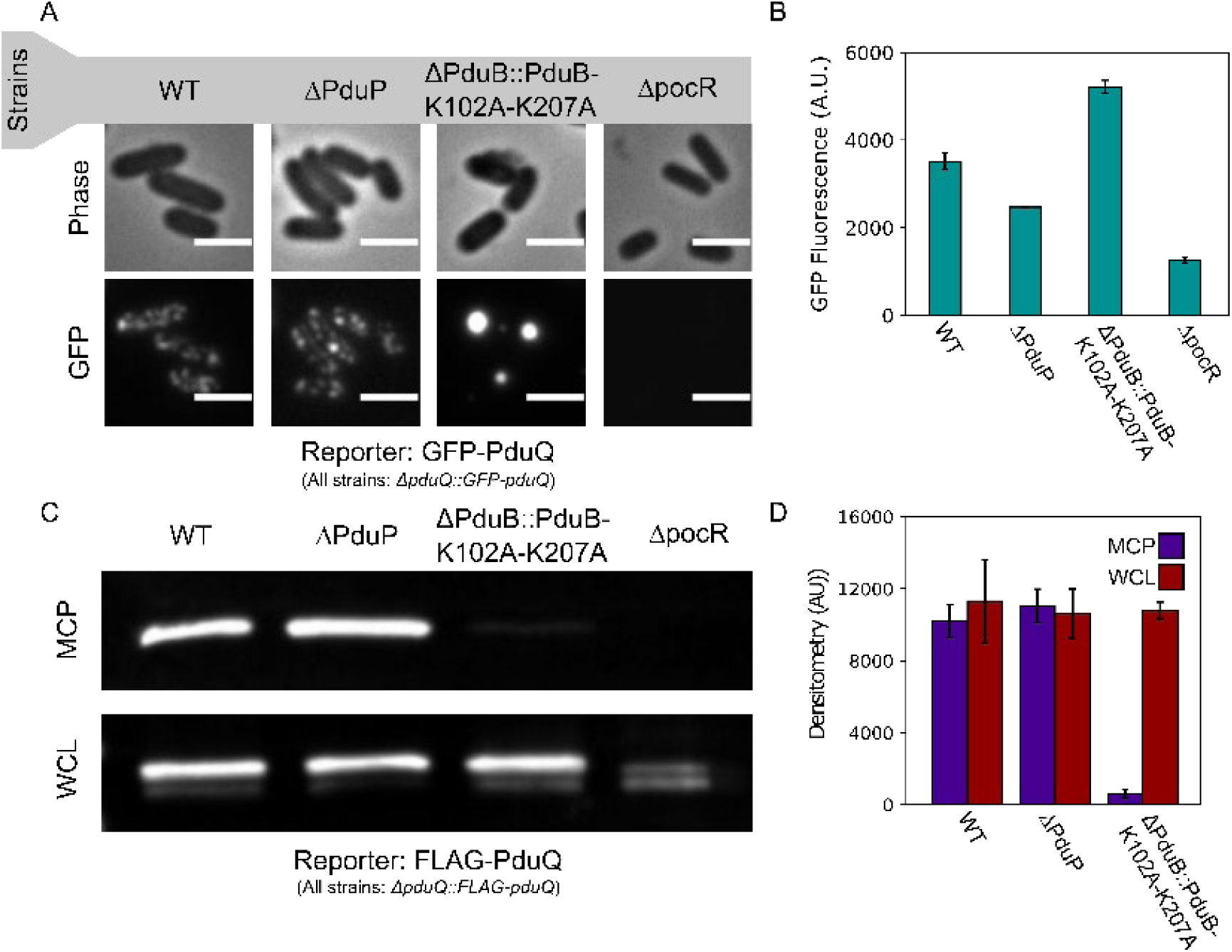
PduQ is encapsulated in the Pdu MCP in the absence of PduP. PduQ reporters were expressed off of the genome at the pduQ locus to mimic the expression and encapsulation of PduQ in the Pdu MCPs used in in vitro reactions. **(A)** Phase contrast and fluorescence micrographs of S. enterica cells expressing GFP-PduQ from the pduQ locus (all ΔpduQ::GFP-pduQ) in different genetic backgrounds (WT, ΔPduP, ΔPduB::PduB-K102A-K207A, and ΔpocR). Bright fluorescent puncta distributed through the cell (e.g., not localized to the pole) are indicative of well-formed MCPs. Scale bars are 2 µm. **(B)** GFP expression was measured via fluorescence to determine if encapsulation efficiency correlated with expression. Fluorescence of GFP-PduQ at the pduQ locus (all ΔpduQ::GFP-pduQ) was measured and normalized to cell density. Error bars indicate standard deviation over three biological replicates. **(C)** Representative anti-FLAG western blot of purified MCPs and whole cell lysate (WCL) samples strains expressing FLAG-PduQ from the pduQ locus (all ΔpduQ::FLAG-pduQ) in different genetic backgrounds (WT, ΔPduP, ΔPduB::PduB-K102A-K207A, and ΔpocR). **(D)** Densitometry of MCP and WCL samples on anti-FLAG western blots of purified MCPs and whole cell lysate samples strains expressing FLAG-PduQ from the pduQ locus (all ΔpduQ::FLAG-pduQ). MCP and WCL densitometry were compared to assess the correlation between encapsulation and expression. Error bars indicate standard deviation across three biological replicates.

**Supplemental Table 2:**
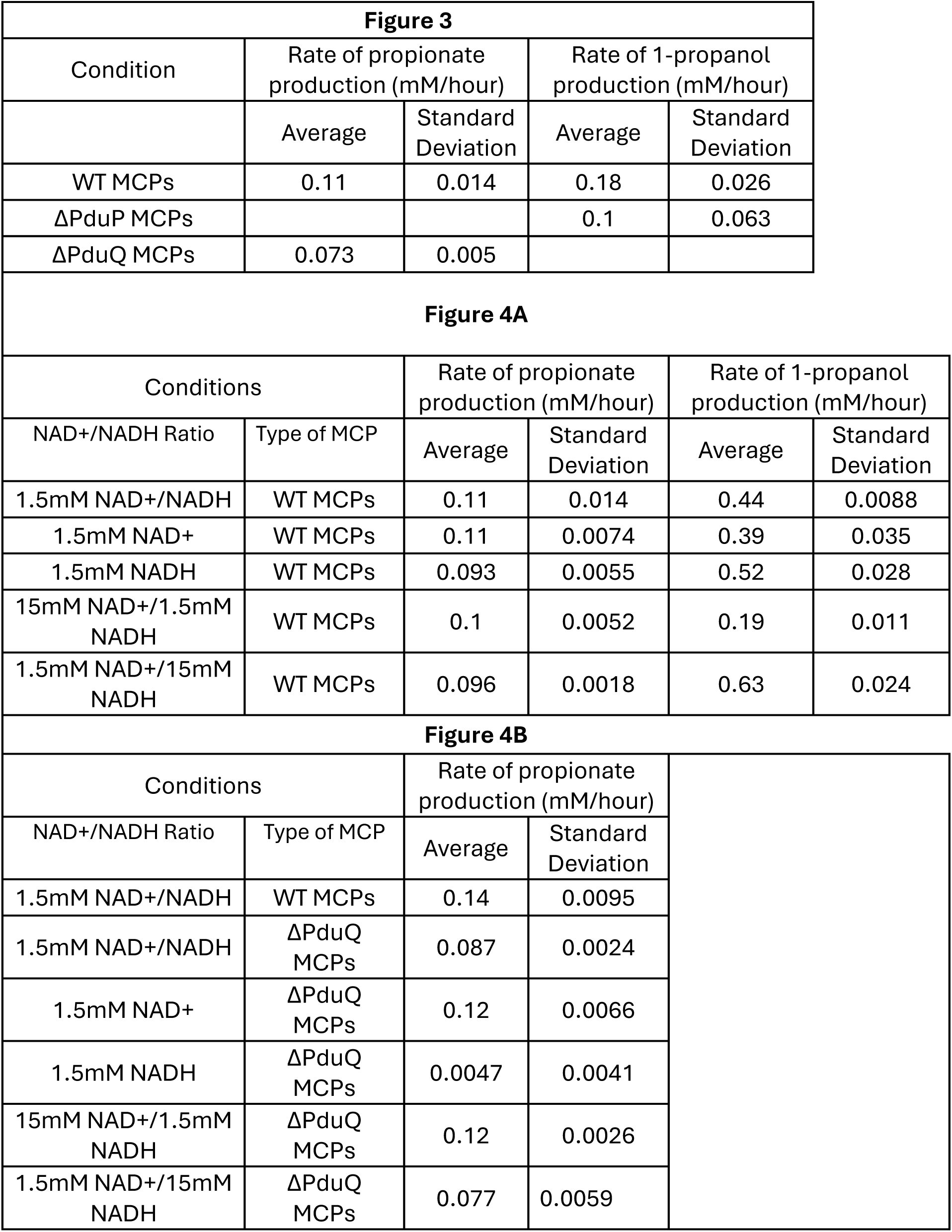
Rates of propionate and 1-propanol production for Figures 3 and 4 from 0 – 6 hours.

**Supplemental Figure 5:**
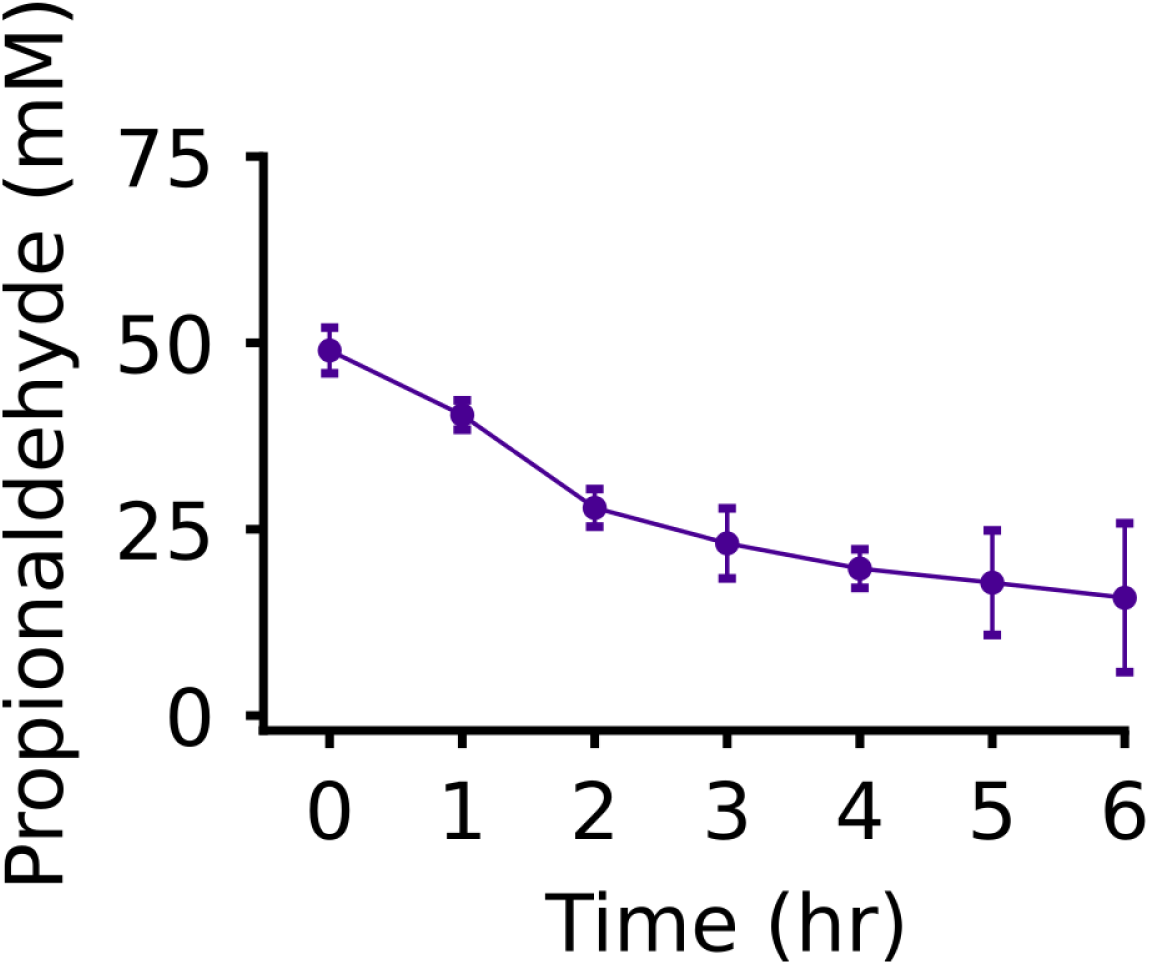
Propionaldehyde evaporates in the *in vitro* assay. 50 mM propionaldehyde was added to *in vitro* reactions lacking Pdu MCPs. The reactions were incubated at 30 °C for 6 hours, and propionaldehyde was measured every hour. Error bars indicated one standard deviation above and below across three technical replicates.

**Supplemental Figure 6:**
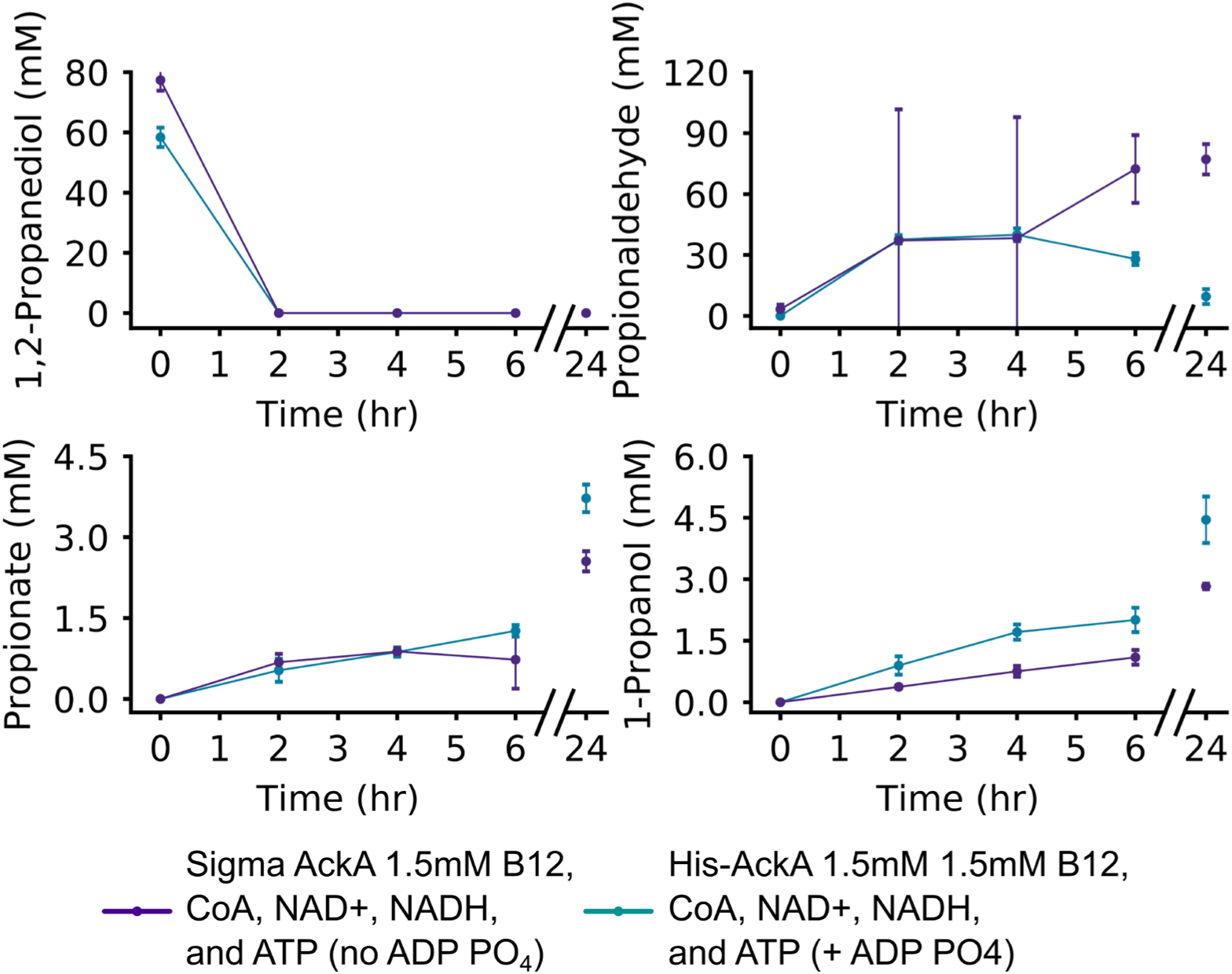
Purified His-AckA performs better than purchased AckA. In vitro measurement of metabolites with WT MCPs with purified His-AckA or AckA purchased (Sigma) added to the reaction. Error bars indicate one standard deviation above and below three technical replicates.

