## Supplemental Information for "Evaluation of Bacterial Microcompartment Cofactor Recycling and Permeability with a Model Guided *In Vitro* Assay"

*Supplemental Table 1: A) Table of strains used in this study. B) Table of primers used in this study.*

A

| **Strain number** | **Strain species** | **Genome** | **Plasmids** | **Description** |
|---|---|---|---|---|
| CHAS063 | *Salmonella enterica* serovar Typhimurium LT2 | WT | - | WT LT2 strain |
| CMJS374 | *Salmonella enterica* serovar Typhimurium LT2 | PduP::cat/sacB | pSIM6 | PduP::cat/sacB |
| TMDS069 | *Salmonella enterica* serovar Typhimurium LT2 | pduQ(PduS RBS)::cat/sacB | pSIM6 | pduQ(PduS RBS)::cat/sacB |
| TMDS150 + PduQ::catsacB | *Salmonella enterica* serovar Typhimurium LT2 | ΔPduP/pduQ(PduS RBS)::cat/sacB | pSIM6 | ΔPduP/pduQ(PduS RBS)::cat/sacB |
| CHAS079 | *Salmonella enterica* serovar Typhimurium LT2 | ΔPduQ | - | ΔPduQ |
| CHAS080 | *Salmonella enterica* serovar Typhimurium LT2 | ΔPduPQ | - | ΔPduPQ |
| TMDS150 | *Salmonella enterica* serovar | ΔPduP | - | ΔPduP |

| | | | | |
|---|---|---|---|---|
| | Typhimurium LT2 | | | |
| CHAS095 | *Salmonella enterica* serovar Typhimurium LT2 | ΔPduGH::catsacB | psim6 | ΔPduGH::catsacB |
| CHAS098 | *Salmonella enterica* serovar Typhimurium LT2 | ΔPduGH | - | ΔPduGH |
| sBJP085 | *Salmonella enterica* serovar Typhimurium LT2 | ΔPduP/PduQ::catsacB | psim6 | ΔPduP/PduQ::catsacB |
| sBJP086 | *Salmonella enterica* serovar Typhimurium LT2 | ΔPduB::PduB-K102A-K207A/PduQ::catsacB | psim6 | ΔPduB::PduB-K102A-K207A/PduQ::catsacB |
| sBJP087 | *Salmonella enterica* serovar Typhimurium LT2 | ΔPduA ΔPduJ/PduQ::catsacB | psim6 | ΔPduA ΔPduJ/PduQ::catsacB |
| sBJP088 | *Salmonella enterica* serovar Typhimurium LT2 | ΔpocR/PduQ::catsacB | psim6 | ΔpocR/PduQ::catsacB |
| sBJP095 | *Salmonella enterica* serovar Typhimurium LT2 | PduQ::FLAG-PduQ | - | PduQ::FLAG-PduQ |
| sBJP096 | *Salmonella enterica* serovar Typhimurium LT2 | ΔPduP/PduQ::FLAG-PduQ | - | ΔPduP/PduQ::FLAG-PduQ |

| | | | | |
|---|---|---|---|---|
| sBJP097 | *Salmonella enterica* serovar Typhimurium LT2 | ΔpduB::PduB-K102A-K207A/PduQ::FLAG-PduQ | - | ΔpduB::PduB-K102A-K207A/PduQ::FLAG-PduQ |
| sBJP098 | *Salmonella enterica* serovar Typhimurium LT2 | ΔPduA ΔPduJ/PduQ::FLAG-PduQ | - | ΔPduA ΔPduJ/PduQ::FLAG-PduQ |
| sBJP099 | *Salmonella enterica* serovar Typhimurium LT2 | ΔpocR/PduQ::FLAG-PduQ | - | ΔpocR/PduQ::FLAG-PduQ |
| sBJP100 | *Salmonella enterica* serovar Typhimurium LT2 | PduQ::GFP-PduQ | - | PduQ::GFP-PduQ |
| sBJP101 | *Salmonella enterica* serovar Typhimurium LT2 | ΔPduP/PduQ::GFP-PduQ | - | ΔPduP/PduQ::GFP-PduQ |
| sBJP102 | *Salmonella enterica* serovar Typhimurium LT2 | ΔpduB::PduB-K102A-K207A/PduQ::GFP-PduQ | - | ΔpduB::PduB-K102A-K207A/PduQ::GFP-PduQ |
| sBJP103 | *Salmonella enterica* serovar Typhimurium LT2 | ΔPduA ΔPduJ/PduQ::GFP-PduQ | - | ΔPduA ΔPduJ/PduQ::GFP-PduQ |
| sBJP104 | *Salmonella enterica* serovar Typhimurium LT2 | ΔpocR/PduQ::GFP-PduQ | - | ΔpocR/PduQ::GFP-PduQ |
| sBJP115 | *Escherichia coli* DH10b | WT | pBJP077 | pBJP077_pET22b_HisNAckA |
| sBJP117 | *Escherichia coli* BL21 DE3 | WT | pBJP077 | pBJP077_pET22b_HisNAckA |

6 B

| Primer | sequencinguence | Description |
|---|---|---|
| TMDP141 | AACCGCGTTTGTAGTGAGAAGGTATTCATCGCGA CCTCAG ttagcgaatagaaaagccgttggtcagtacaatcagggtttcgagt tcagaagtattcat | Linear DNA used to knockout PduP |
| TMDP086 | tcatttacagggaaaagtggtcacc | PduP KO sequencing (internal) f |
| TMDP087 | tgcgccagaaagccatcg | PduP KO sequencing (internal) r |
| TMDP088 | tgtgacctggcggatgc | PduP KO sequencing (external) f |
| TMDP089 | tgcagagcctgcatttgc | PduP KO sequencing (external) r |
| TMDP152 | CCCGTTCCCGGCGCTGCGTACTGACCAACGGCT TTTCTATTCGCTAACTGAGGTCGCGatgaataccttctc actacaaacgcgg | PduQ KO fwd primer |
| TMDP153 | AAGGGACATTTCAACGCTGTTGATGGCGGTGCTC ATAGCAGTTCCTCCAGCAGTTCccgaatcaaccgcgtt tgtagtgagaagg | PduQ KO rev primer |
| TMDP097 | gaaggctttaccaccttcactatcg | PduQ KO internal sequencing f |
| TMDP098 | cctgcgcctgtagtttgacg | PduQ KO internal sequencing r |
| TMDP099 | cataccgccattatgcactcg | PduQ KO sequencing (external) f |
| TMDP100 | agggctatcaccccttcg | PduQ KO sequencing (external) r |
| CHA195 | atgcgatatatagctggcattgacatcggtgggatcccctttcggg atctccatgcttaa | PduGH_KO fwd primer |
| CHA196 | ttaagcatggagatcccgaaaggggatcccaccgatgtcaatgc cagctatatatcgcat | PduGH_KO rev primer |
| CHA197 | cttcgaacgcgccgccg | PduGHKOsequencingf_ amp |
| CHA198 | ccacgacgcttgccgctg | PduGHKOsequencingr_ amp |
| CHA199 | agctttcgttcgtgaagcggcag | PduGHKOsequencingf |
| oBJP176 | tcggcgctaagggacatttcaacgctgttgatggcggtgctcatTT Atagcagttcctccagcagt | Reverse primer to add PduQ homology to PduQ |
| oBJP177 | tactgaccaacggcttttctattcgctaactgaggtcgcgATGGA TTATAAAGATGACGATgacaag | FWD primer to add PduQ locus homology to N terminally tagged FLAG |

| | | |
|---|---|---|
| oBJP178 | tactgaccaacggcttttctattcgctaactgaggtcgcgATGAg taaaggagaagaacttttca | FWD primer to add PduQ locus homology to N terminally tagged GFP |

9

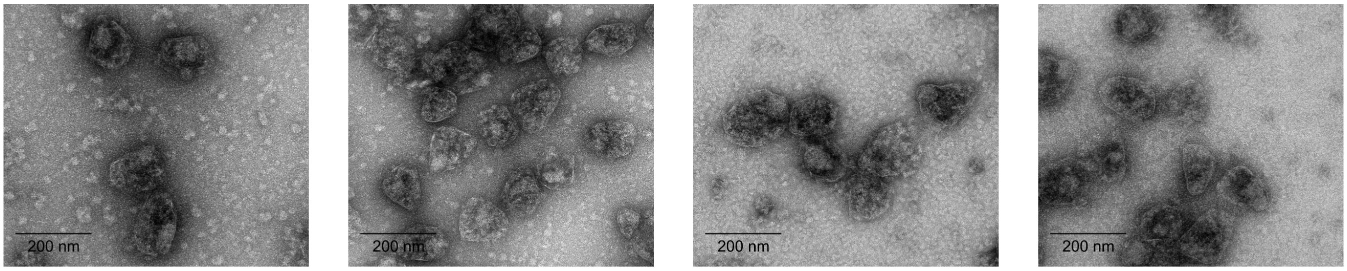


WT ΔPduP ΔPduQ ΔPduPQ

**Supplemental Figure 1: TEM images of purified Pdu MCPs-** Transmission electron micrographs of purified WT Pdu MCPs, ΔPduP Pdu MCPs, ΔPduQ Pdu MCPs, and ΔPduPQ Pdu MCPs. Scale bars are 200 nm.

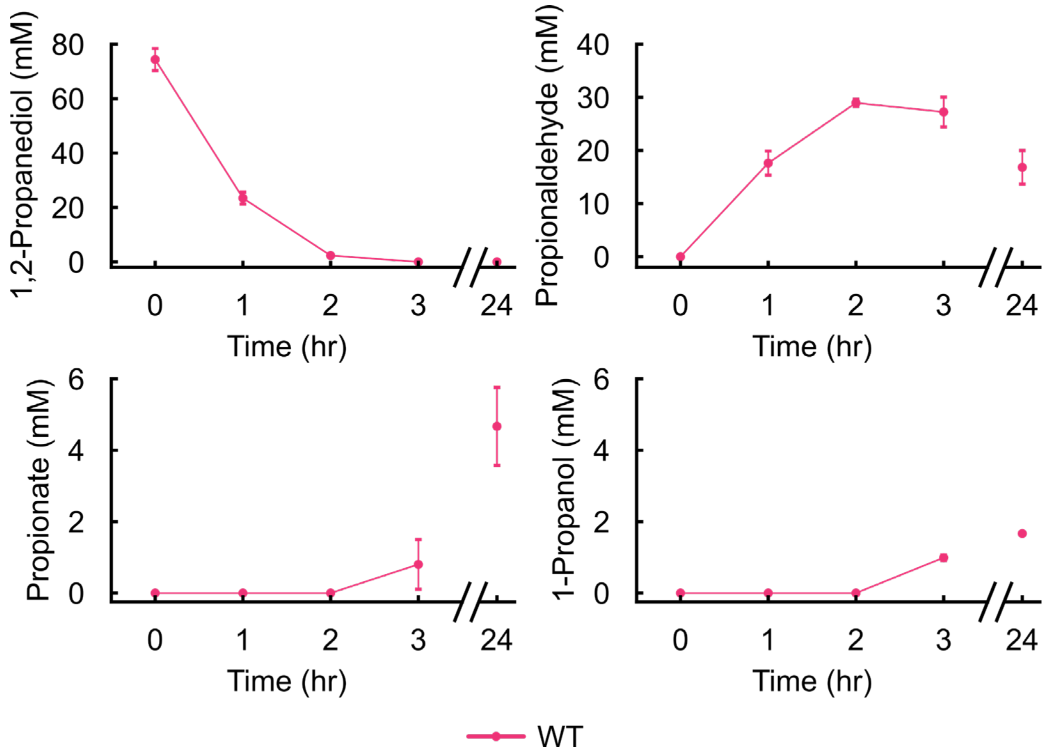


**Supplemental Figure 2: Combining purified Pdu MCPs, 1,2-PD, and the Pdu pathway cofactors results in full pathway activation in vitro –** WT Pdu MCPs were added to reaction buffer containing the following: 1,2-PD, $B_{12}$, CoA, NAD+, NADH, and ATP. Concentrations of the metabolites 1,2-PD, propionaldehyde, propionate, and 1-propanol were measured at 0, 1, 2, 3, and 24 hours. Error bars indicate one standard deviation of three technical replicates.

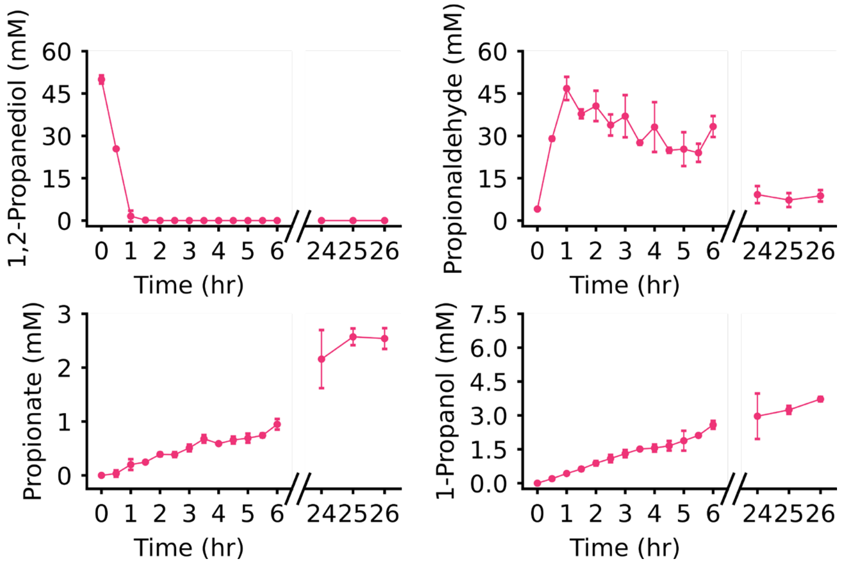


**Supplemental Figure 3: Steady state is reached at 24 hours –** WT Pdu MCPs were assayed with measurements taken from 0-6 hours and from 24-26 hours to show steady state at 24 hours. WT Pdu MCP metabolite data from 0-6 hours and 24 hours was also displayed in Figure 4B. Error bars indicated one standard deviation above and below across three technical replicates.

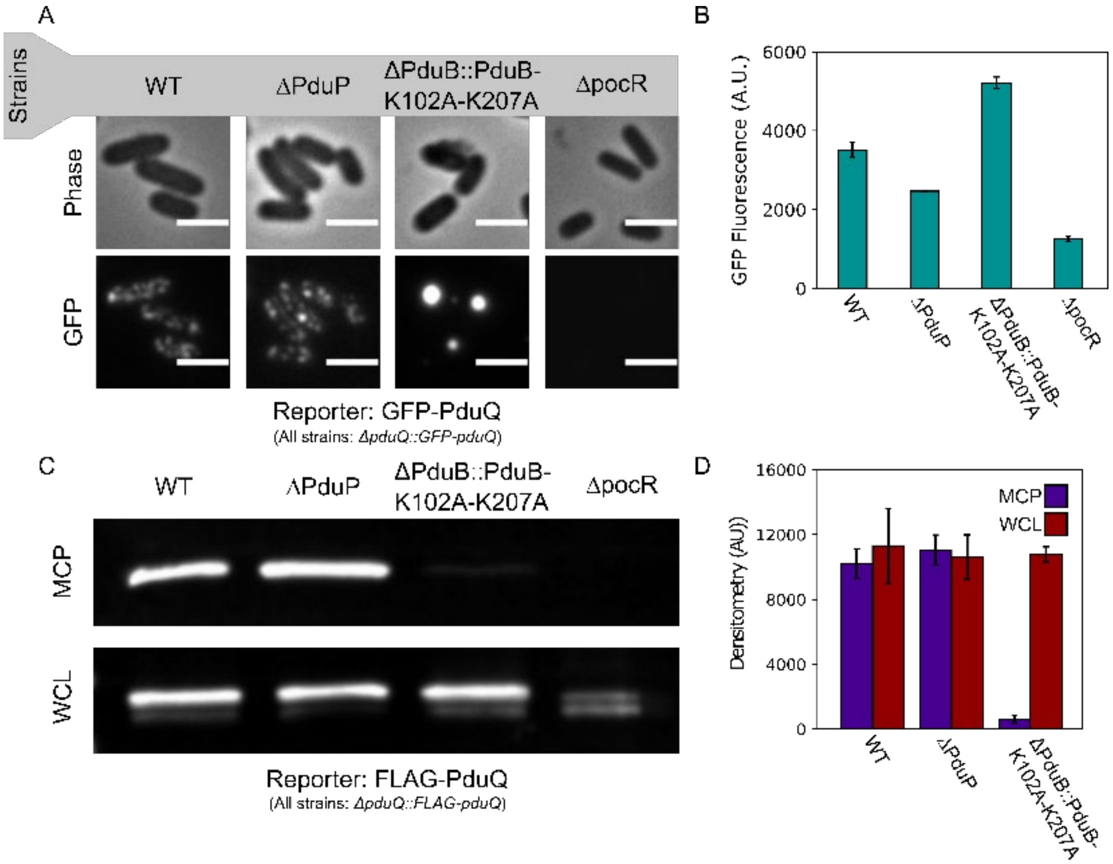


**Supplemental Figure 4: PduQ is encapsulated in the Pdu MCP in the absence of PduP -** PduQ reporters were expressed off of the genome at the pduQ locus to mimic the expression and encapsulation of PduQ in the Pdu MCPs used in in vitro reactions. **(A)** Phase contrast and fluorescence micrographs of S. enterica cells expressing GFP-PduQ from the pduQ locus (all ΔpduQ::GFP-pduQ) in different genetic backgrounds (WT, ΔPduP, ΔPduB::PduB-K102A-K207A, and ΔpocR). Bright fluorescent puncta distributed through the cell (e.g., not localized to the pole) are indicative of well-formed MCPs. Scale bars are 2 μm. **(B)** GFP expression was measured via fluorescence to determine if encapsulation efficiency correlated with expression. Fluorescence of GFP-PduQ at the pduQ locus (all ΔpduQ::GFP-pduQ) was measured and normalized to cell density. Error bars indicate standard deviation over three biological replicates. **(C)** Representative anti-FLAG western blot of purified MCPs and whole cell lysate (WCL) samples strains expressing FLAG-PduQ from the pduQ locus (all ΔpduQ::FLAG-pduQ) in different genetic backgrounds (WT, ΔPduP, ΔPduB::PduB-K102A-K207A, and ΔpocR). **(D)** Densitometry of MCP and WCL samples on anti-FLAG western blots of purified MCPs and whole cell lysate samples strains expressing FLAG-PduQ from the pduQ locus (all ΔpduQ::FLAG-pduQ). MCP and WCL densitometry were compared to assess the correlation between encapsulation and expression. Error bars indicate standard deviation across three biological replicates.

*Supplemental Table 2: Rates of propionate and 1-propanol production for Figures 3 and 4 from 0 – 6 hours*

**Figure 3**

| Condition | Rate of propionate production (mM/hour) | | Rate of 1-propanol production (mM/hour) | |
|---|---|---|---|---|
| | Average | Standard Deviation | Average | Standard Deviation |
| WT MCPs | 0.11 | 0.014 | 0.18 | 0.026 |
| ΔPduP MCPs | | | 0.1 | 0.063 |
| ΔPduQ MCPs | 0.073 | 0.005 | | |

**Figure 4A**

| Conditions | | Rate of propionate production (mM/hour) | | Rate of 1-propanol production (mM/hour) | |
|---|---|---|---|---|---|
| NAD+/NADH Ratio | Type of MCP | Average | Standard Deviation | Average | Standard Deviation |
| 1.5mM NAD+/NADH | WT MCPs | 0.11 | 0.014 | 0.44 | 0.0088 |
| 1.5mM NAD+ | WT MCPs | 0.11 | 0.0074 | 0.39 | 0.035 |
| 1.5mM NADH | WT MCPs | 0.093 | 0.0055 | 0.52 | 0.028 |
| 15mM NAD+/1.5mM NADH | WT MCPs | 0.1 | 0.0052 | 0.19 | 0.011 |
| 1.5mM NAD+/15mM NADH | WT MCPs | 0.096 | 0.0018 | 0.63 | 0.024 |

**Figure 4B**

| Conditions | | Rate of propionate production (mM/hour) | |
|---|---|---|---|
| NAD+/NADH Ratio | Type of MCP | Average | Standard Deviation |
| 1.5mM NAD+/NADH | WT MCPs | 0.14 | 0.0095 |
| 1.5mM NAD+/NADH | ΔPduQ MCPs | 0.087 | 0.0024 |
| 1.5mM NAD+ | ΔPduQ MCPs | 0.12 | 0.0066 |
| 1.5mM NADH | ΔPduQ MCPs | 0.0047 | 0.0041 |
| 15mM NAD+/1.5mM NADH | ΔPduQ MCPs | 0.12 | 0.0026 |
| 1.5mM NAD+/15mM NADH | ΔPduQ MCPs | 0.077 | 0.0059 |

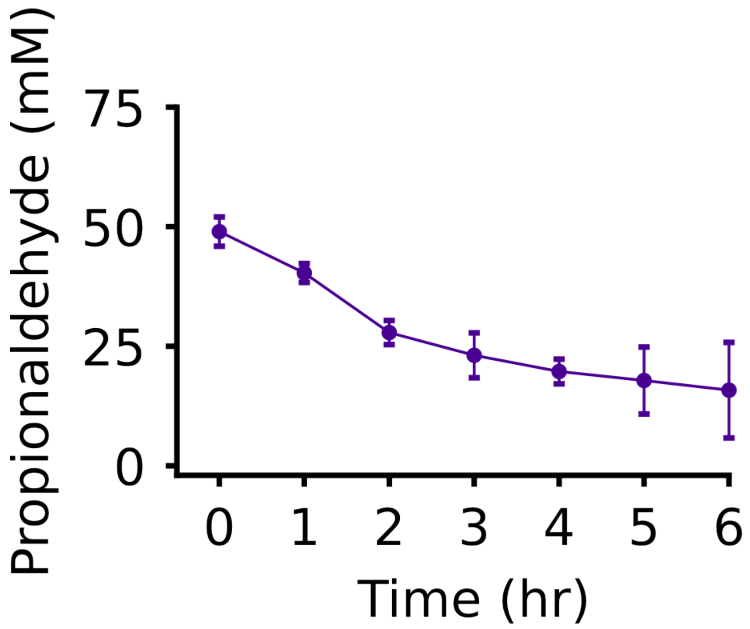


**Supplemental Figure 5: Propionaldehyde evaporates in the *in vitro* assay –** 50 mM propionaldehyde was added to *in vitro* reactions lacking Pdu MCPs. The reactions were incubated at 30 °C for 6 hours, and propionaldehyde was measured every hour. Error bars indicated one standard deviation above and below across three technical replicates.

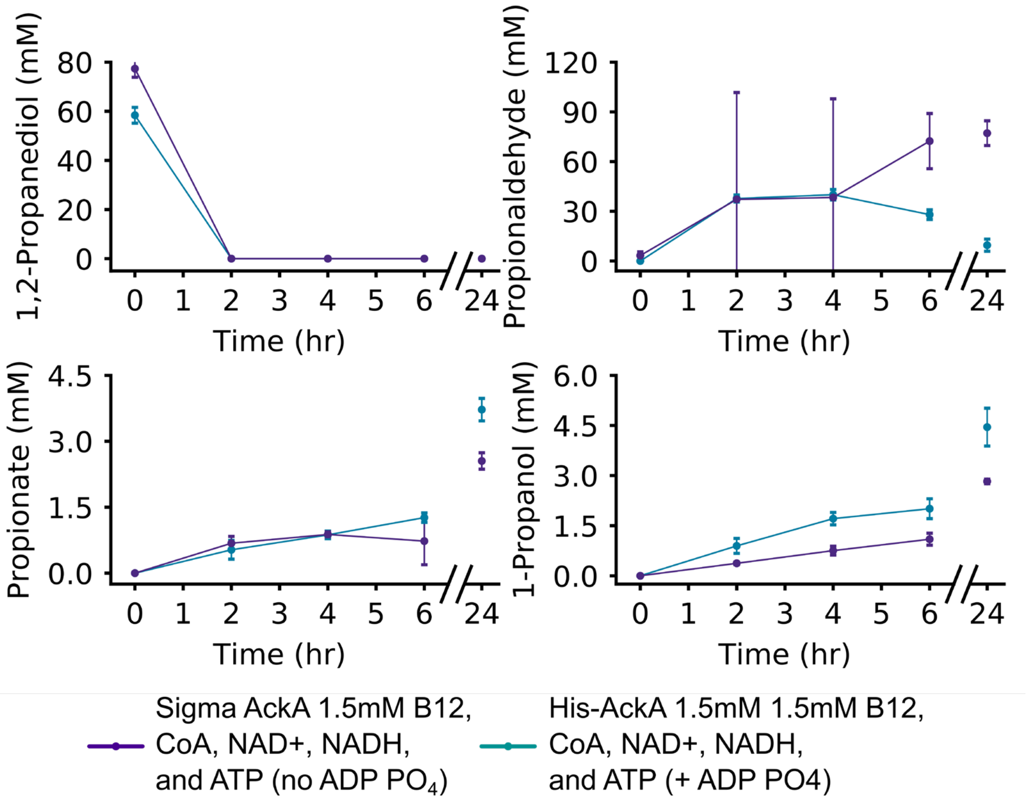


**Supplemental Figure 6: Purified His-AckA performs better than purchased AckA -** In vitro measurement of metabolites with WT MCPs with purified His-AckA or AckA purchased (Sigma) added to the reaction. Error bars indicate one standard deviation above and below three technical replicates.
